# Trimethylamine N-Oxide (TMAO) links peripheral insulin resistance and cognitive deficiencies in a senescence accelerated mouse model

**DOI:** 10.1101/2021.11.17.469001

**Authors:** Manuel H. Janeiro, Elena Puerta, María Lanz, Fermin I. Milagro, María J. Ramírez, Maite Solas

## Abstract

It has been established that ageing is the major risk factor for cognitive deficiency or neurodegenerative diseases such as Alzheimer’s disease (AD) and it is becoming increasingly evident that insulin resistance is another factor. Biological plausibility for a link between insulin resistance and dementia is relevant for understanding disease etiology, and to form bases for prevention efforts to decrease disease burden.

The dysfunction of the insulin signaling system and glucose metabolism has been proposed to be responsible for brain aging. Normal insulin signaling in the brain is required to mediate growth, metabolic functions, and the survival of neurons and glia. Insulin receptors are densely expressed in the olfactory bulb, the cerebral cortex and the hippocampus and regulate neurotransmitter release and receptor recruitment. In normal elderly individuals, reduced glucose tolerance and decreased insulin levels in the aged brain are typically observed. Furthermore, insulin signaling is aberrantly activated in the AD brain, leading to non-responsive insulin receptor signaling.

The senescence accelerated mouse (SAMP8) mouse was one of the accelerated senescence strains that spontaneously developed from breeding pairs of the AKR/J series. The SAMP8 mouse develops early learning and memory deficits (between 6 and 8 months) together with other characteristics similar to those seen in Alzheimer’s disease. The present project proposes the investigation of the missing link between aging, insulin resistance and dementia.

Peripheral but not central insulin resistance was found in SAMP8 mice accompanied by cognitive deficiencies. Furthermore, a marked peripheral inflammatory state (i.e. significantly higher adipose tissue TNF-α and IL6 levels) were observed in SAMP8 mice, followed by neuroinflammation that could be due to a higher cytokine leaking into the brain across a aging-disrupted BBB. Moreover, aging-induced gut dysbiosis produces higher TMAO that could also contribute to the peripheral and central inflammatory tone as well as to the cognitive deficiencies observed in SAMP8 mice. All those alterations were reversed by DMB, a treatment inhibits the transformation of choline, carnitine and crotonobetaine, decreaseing TMAO levels.

The ever-increasing incidence of neurodegenerative diseases not only limits the life quality of the affected individuals and their families but also poses an enormous demand on the societies. Thus, it is instrumental to pursue novel promising approaches to prevent and treat it at the highest possible speed to rapidly translate them to clinical practice. From this point of view, data obtained from this project will be instrumental to validate the principle approach of microbial dysbiosis and increased TMAO secretion as a key link between aging, insulin resistance and dementia. Collectively, the proposed experiments ideally integrate the aim to promote a novel approach to improve the lives of those suffering from cognitive disturbances.

## Introduction

Aging is a natural process which is often related to inflammation and the progressive impairment of several physiological functions including metabolic homeostasis, leading to insulin resistance (IR) (Fang et al. 2018).

Recent studies have hypothesized that peripheral metabolic dysregulation may play an important role in cognitive decline, suggesting similar mechanisms between neuronal and peripheral IR (Frazier et al. 2019). High peripheral insulin levels have been related with poor cognitive function in aged patients (Kuusisto et al. 1993; Stolk et al. 1997). Moreover, a chronic elevation of insulin levels was also linked to a major decline in performance in healthy aged people, showing that peripheral IR and a prediabetic state were also associated to worse performances (Burns et al. 2012). However, the molecular mechanisms underlying this crosstalk are still elusive, as well as how central and peripheral insulin signaling operate in cognitive decline (Biessels and Despa, 2018).

Accumulating evidence has shown that crosstalk between insulin signaling and low-grade inflammatory tone could be one of the main processes leading to the initiation of neuronal insulin resistance (Thaler et al., 2013; Velloso et al., 2011; Konner et al., 2011). Hence, activation of inflammatory cascades within the brain, either secondary to elevated saturated free fatty acid or associated with increased release of proinflammatory cytokines, impairs insulin signaling directly (Velloso et al., 2011; Konner et al., 2011).I ncreased levels of pro-inflammatory cytokines, particularly tumor necrosis factor alpha (TNF-α) activates c-Jun terminal kinase (JNK), resulting in insulin receptor substrate 1 (IRS1) suppression by increasing the inhibitory phosphorylation and decreasing activating phosphorylation (Hotamisligil et al., 1993; Rui et al., 2001; Pedersen et al., 2003). Interestingly, local inflammation is also a common feature of brain aging. Glial cells, often exhibit an activated state in the aged brain that is characterized by the acquisition of an ameboid morphology and the production of pro-inflammatory cytokines including interleukin 1b (IL-1b), IL-6, and TNF-α (Cribbs et al., 2012; Norden and Godbout, 2013). Noteworthy, BBB integrity can be disrupted by neuroinflammation associated with many age-related disorders as Alzheimer’s disease (Cai et al., 2018). Thus, aberrant amounts of plasma pro-inflammatory cytokines may enter the brain via the BBB and cause inflammation by altering microglial (Erny et al. 2015) and astrocyte activation (Rothhammer et al. 2016) and subsequent brain cytokine release. Therefore, insulin resistance and neuroinflammation are interconnected pathological features and both are directly or indirectly considered to be two major culprits in cognitive disturbances (Ferreira et al., 2014).

On the other hand, aging is also characterized by changes in gut microbiota composition decreasing its diversity, what could play a key role in the development of several disturbances such us IR and even neuropathologies (Janeiro et al., 2021). Dysbiosis is able to induce reduced expression of tight junction proteins in the gut, inducing intestinal inflammation and a major permeability, what favours the passage of gut metabolites to the circulatory system (Janeiro et al., 2021). Among these microbial metabolites, trimethylamine N-oxide (TMAO) has been recently shown to exacerbate IR (Chen et al. 2019; Gao et al. 2015; Oellgaard et al. 2017) and it has been also related to inflammation and cognitive decline (Janeiro et al. 2018). Moreover, mice fed a Western diet, a risk factor for insulin resistance, have greater plasma TMAO concentrations and show a higher expression of pro-inflammatory cytokines, such as tumor necrosis factor-α (TNF-α) and interleukin-1β (IL-1β) and a decrease in the expression of anti-inflammatory cytokines (IL-10). The treatment with 3,3-dimethyl-1-butanol (DMB), that inhibits the choline TMA lyase enzyme (one of the key enzymes for TMAO synthesis), prevented all those outcomes and lowered plasma TMAO levels (Chen et al., 2017).

To elucidate the basic mechanisms of age-related changes and develop effective drugs for the prevention of age-related diseases, the establishment of appropriate animal models with human-like characteristics is essential. SAM (senescence-accelerated mouse) has already been established as a mouse model for accelerated aging. It is actually made up of a group of related inbred mice that include nine short-lived animal strains susceptible to accelerated senescence (SAMP) and three long-lived mice resistant to accelerated senescence (SAMR). Specifically, in the present work, SAMP8 strain has been chosen as it shows relatively specific age-associated phenotypic pathologies such as a shortened life span and early manifestation of senescence. Furthermore, the SAMP8 mouse develops early learning and memory deficits (between 8 and 10 months) together with other characteristics similar to those seen in Alzheimer’s disease, such as, elevated amyloid-beta burden (Takemura et al., 1993; Fukunari et al., 1994; Morley et al., 2000), hyperphosphorylation of tau (Canudas et al., 2005), increased alpha synuclein (Caballero et al., 2008), an increase in presinilin (Kumar et al., 2009), increased oxidative damage (Butterfield et al., 2005), decreased choline acetyl transferase activity (Strong et al., 2003), increased glutamate (Kitamura et al., 1992), altered NMDA function (Nomura et al., 1997) and increased neuronal nitric oxide synthase (Ali et al., 2009).

Hence, in this study, the plausible link between peripheral insulin resistance and cognitive decline has been investigated, with a focus on inflammation and gut dysbiosis as persuasive linking mechanisms.

## Material and Methods

### Animals

All the experiments were carried out in a total of 66 mice which were divided in 6 subgroups: SAMR1 2 months-old (n=11), SAMP8 2 month-old (n=8), SAMR1 6 month-old (n=12), SAMP8 6 month-old (n=10), SAMR1 10 month-old (n=17) and SAMP8 10 month-old (n=8). Number of animals was dependant on availability. Animals were housed in a temperature- (21 ± 1 °C) and humidity- (55 ± 5%) controlled room on a 12-h light/dark cycle.

For DMB (3,3-dimethyl-1-butanol) treatment, eighteen SAMR1 and eighteen SAMP8 mice aged 6 months were randomly divided into four experimental groups as follows (n=9 per group): SAMR1 control group (R1-C), SAMR1 mice treated with 1% (vol/vol) DMB group (R1-DMB), SAMP8 mice control group (P8-C), SAMP8 mice treated with 1% (vol/vol) DMB group (P8-DMB). DMB (TCIAD1333, VWR, TCI Europe) was given in drinking water for 9 weeks and renewed every week.

Animal handling and breeding was conducted in accordance with the principles of laboratory animal care as detailed in the European Communities Council Directive (2003/65/EC), Spanish legislation (Real Decreto 1201/2005) and approved by the Ethics Committee of the University of Navarra. In addition, every effort was made to minimize the number of animals used and any possible suffering.

### Body composition assessment

Body composition assessment was made using quantitative magnetic resonance (QMR). Scans were performed by placing animals into a transparent plastic cylinder (1.5 mm thick, 4.7 cm diameter), with a smaller plastic cylinder inserted into the big one to limit mouse movement. While in the tube, animals were briefly subjected to a low-intensity (0.05 Tesla) electromagnetic field to measure fat and lean mass.

### Glucose- and insulin-tolerance tests

GTT was performed after mice underwent a fasting period of 6 h. Glucose concentrations in blood were measured after the fasting period (0 min), then each mouse received an intraperitoneal injection of 20% glucose (10 ml per kg body weight) and glucose concentrations in blood were measured after 15, 30, 60 and 120 min.

ITT was done with mice fed ad libitum. After basal glucose concentrations in blood were measured (0 min), each mouse received an intraperitoneal injection of insulin (0.75 units per kg body weight; Actrapid; Novo Nordisk) and glucose concentrations in blood were measured after 15, 30 and 60 min.

### Behavioural test

Behavioural experiments were conducted between 09:00 h and 13:00 h. To perform these tests animals were randomized.

#### Open field

Locomotor activity was measured for 30 min in an open field (65 × 65 cm^2^, 45 cm height) made of black wood, using a video-tracking system (Ethovision 3.0, Noldus Information Technology B.V., The Netherlands), in a softly illuminated room. Total path length (cm) was analysed.

#### Novel object recognition test (NORT)

The open field consisted of a square divided into four sections (65 cm × 65 cm × 45 cm each) with black walls. On the previous day to the experiment, animals were familiarized with the square for 30 min. The test consists of 3 trials of 5 minutes: sample phase, 1 hour trial and 24 hours trial. During the first trial, two identical objects were placed inside the cubicle, and the mice were allowed to explore. An hour later the second trial took place, in which one object was replaced by another, and exploration was scored for 5 min. Twenty-four hours later the third task took place where once again one object was changed and the exploration time was recorded for 5 min. Results were expressed as percentage of time spent exploring the new object with respect to the total exploration time (discrimination index). It is important to highlight that the exploration was considered complete when the nose of the mouse was oriented within 2 cm of the object.

#### Morris Water Maze

The Morris water maze (MWM) was used to test spatial memory and to evaluate the working and reference memory functions in SAMP8 mice.

The water maze was a circular pool (diameter of 145 cm) filled with water (21–22 °C) and virtually divided into four equal quadrants identified as northeast, northwest, southeast, and southwest.

To test learning capacity, hidden-platform training was conducted with the platform placed in the northeast quadrant 1 cm below the water surface over 8 consecutive days (4 trials/day). Several large visual cues were placed in the room to guide the mice to the hidden platform. Each trial was finished when the mouse reached the platform (escape latency) or after 60 s, whichever came first. Mice failing to reach the platform were guided onto it. After each trial mice remained on the platform for 15 s. To test memory, probe trials were performed at the 4^th^, 7^th^ and last day of the test (9^th^ day). In the probe trials the platform was removed from the pool and mice were allowed to swim for 60 s. The percentage of time spent in the target quadrant was recorded. All trials were monitored by a video camera set above the center of the pool and connected to a video tracking system (Ethovision 3.0; Noldus Information Technology B.V, Wageningen, Netherlands).

### Tissue and blood collection

Mice were killed by decapitation between 09:00–12:00 h. Brains were removed and dissected on ice to obtain the hippocampus and stored at −80 °C.

For immunohistochemistry assays, left hemispheres from 5 mice per group were fixed by immersion in 4% paraformaldehyde in 0.1 M PBS (pH 7.4) for 24 h followed by 20% sucrose solution. Brains were cut into series of 40 μm slides.

Blood was collected, centrifuged at 1250 g (15 min, 4°C), and serum was frozen at −80°C.

### Plasma insulin and glucose levels

Insulin and glucose were measured in plasma samples using the Sensitive Insulin enzyme immunoassay kit (EZRMI-13K, Millipore, Billerica, MA) and Glucose GOD-PAP enzyme immunoassay kit (Roche Diagnostics, Spain) respectively, following the manufacturer’s instructions.

Insulin sensitivity was analysed by assessing the homeostatic model assessment (HOMA) index. HOMA index is expressed as insulin (μU/mL) × glucose (mg/dL) / 405.

### Western Blotting

Assays were performed in hippocampal tissue as described previously (Solas et al. 2010). Samples (50 μg of protein) were separated by electrophoresis on a sodium dodecyl sulphate-polyacrylamide gel (7.5%). Membranes were probed overnight at 4 °C with the corresponding primary antibodies (Table 1). Secondary antibodies conjugated to IRDye 800CW or IRDye 680CW (LI-COR Biosciences, Lincoln, NE, USA) were diluted to 1/15,000 in TBS with 5% BSA. Bands were visualized using Odyssey Infrared Imaging System (LI-COR Biosciences, Lincoln, NE, USA). β-actin was used as internal control. Results were calculated as the percentage of optical density values of the SAMR1 mice.

**Table 1.**
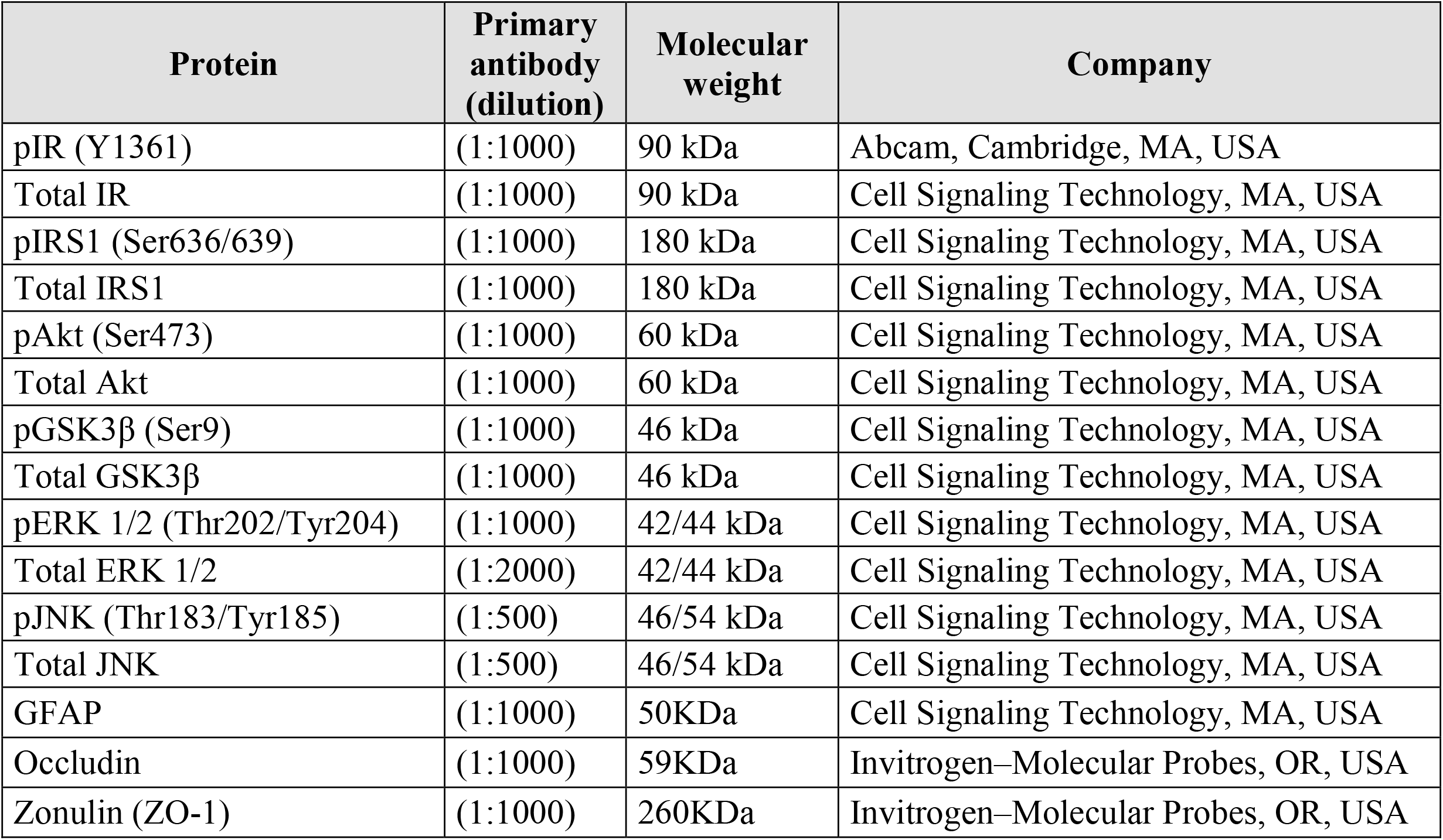
Primary antibodies used for western blot experiments

### Immunofluorescence staining

Serial coronal brain slices (thickness: 40 μm) were cut with a freezing microtome from the frontal cortex till the end of the hippocampus, and were stored in a cryoprotectant solution. Floating tissue sections comprising the hippocampus were processed for immunohistochemistry.

Free-floating brain sections were washed (3 × 10 min) with PBS 0.1 M (pH 7.4) and incubated in blocking solution (PBS containing 0.3% Triton X-100, 0.1% BSA and 2% normal donkey serum) for 2 h at room temperature. Primary and secondary antibodies were diluted in the blocking solution. Sections were incubated with the primary antibody overnight at 4°C, washed with PBS and incubated with the secondary antibody for 2 h at room temperature, protected from light. The primary antibodies used were anti-GFAP (1:250, Cell Signalling Technology, Beverly, MA, USA), anti-fibrin (1: 1000, Dako, Santa Clara, CA, USA), anti-ZO-1 (1: 200, Invitrogen–Molecular Probes, Eugene, OR, USA) and anti-Occludin (1: 200, Invitrogen–Molecular Probes, Eugene, OR, USA). Secondary antibodies used was Alexa Fluor 546 goat anti-mouse (1:200, Invitrogen–Molecular Probes, Eugene, OR, USA) and alexa Fluor 546 goat anti-rabbit (1: 400, Invitrogen–Molecular Probes, Eugene, OR, USA). To visualize brain microvessels, fluorescein-conjugated Lycopersicon esculentum lectin (1: 200, Vector Laboratories, Burlingame, CA, USA) was used and incubated together with the secondary antibody. For better visualization of nuclei, sections were rinsed 10min in the DNA marker TOPRO-3 (Invitrogen–Molecular Probes, Eugene, OR, USA) working concentration 4 μM in PBS, and then washed 2 min in PBS before mounting. To ensure comparable immunostaining, sections were processed together under identical conditions. Fluorescence signals were detected with confocal microscope LSM 510 Meta (Carl Zeiss, Oberkochen, Germany).

Images were randomly taken from three non-adjacent tissue sections per specimen (n=4) and analyzed using NIH-developed ImageJ. To quantify capillary leakage, the levels of extravascular fibrin and IgG were measured as previously described (Bell et al., 2010; 2012; Winkler et al., 2012; 2013). Briefly, the ImageJ Area tool was used to measure the total area of fibrin- and IgG-positive signal and, when it colocalized with the lectin-positive signal, it was subtracted from the total area of leakage. Using this method, a value representing extravascular levels of each plasma-derived protein is obtained. All images were analyzed by a blinded investigator.

### Real time quantitative PCR assay

Total RNA was obtained from white adipose tissue, brain frontal cortex and brain hippocampus, by homogenizing in TRIzol (Thermo Scientific, Rockford, IL, USA) and following standard methods. RNA quantity and quality were evaluated by a Nanodrop ND-1000 spectrophotometer (Thermo Scientific, Rockford, IL, USA). Then, 1 μg of the total RNA of each sample was reverse-transcribed into cDNA using SuperScript III cDNA Synthesis Kit (Thermo Scientific, Rockford, IL, USA). Real-time PCRs were performed using the TaqMan™ Universal Master Mix (Thermo Fisher scientific, MA, USA) in an CFX384 Touch real-time PCR Detection System (Bio-Rad, Hercules, CA, USA). As housekeeping gene, Glyceraldehyde 3-phosphate dehydrogenase (GAPDH) was chosen. The specific primers used for cDNA amplification were interleukin 6 (Il-6) (Mm00446190_m1) and tumour necrosis factor-alpha (Tnf-α) (Mm00443258_m1). Samples were analysed by a double delta CT (ΔΔCT) method. Relative transcription levels (2−ΔΔCt) were expressed as a mean ± standard error of the mean.

### Fecal Sample Collection, Bioinformatics and Metagenomic Data

Fecal samples were collected using OMNIgene.GUT kits from DNA Genotek (Ottawa, ON, Canada), that is able to stabilize gut microbiome composition and can be kept at room temperature. CIMA LAB Diagnostics (University of Navarra, Pamplona, Spain) was responsible of bacterial DNA sequencing. In order to characterize the phylogeny and taxonomy of the microbial samples the bacterial 16S RNA gene was sequenced. This gene has approximately 1500 base pairs and contains 9 variable regions in between the conserved regions. For the 16S RNA gene analysis, the V3 and V4 hypervariable regions of the gene were amplified followed by the sequencing, which allowed the assignation up to the species level.

The analyses were conducted using the the Illumina MiSeq equipment, following this protocol: the V3 and V4 regions of the 16S gene are amplified in two PCR reactions that are carried out in a thermocycler, creating an amplicon of approximately 460 base pairs. This requires the use of the 16S-F and 16S-R specific primers (16S Forward Primer =5 0 TCGTCGGCAGCGTCAGATGTGTATAAGAGACAGCCTACGGGNGGCWGCAG; 16S Reverse Primer = 5 0 GTCTCGTGGGCTCGGAGATGTGTATAAGAGACAGGACTACHVGGGTATCT AATCC). The protocol for the first PCR reaction was the following: 95 °C for 3 minutes, and 25 cycles of 95 °C for 30 seconds, 55 °C for 30 seconds, 72 °C for 30 seconds, and finally, 72 °C for 5 minutes, to later keep refrigerated at 4 °C. After the cleansing process, 5μl were extracted from the first PCR reaction sample to use for the second PCR reaction. For the second PCR reaction the employed protocol was 95 °C for 3 minutes, and 8 cycles of 95 °C for 30 seconds, 55 °C for 30 seconds, 72 °C for 30 seconds, and finally, 72 °C for 5 minutes, to later keep refrigerated at 4 °C. After each PCR reaction, a cleansing process was carried out to clear the sample from primers. Then, in order to sequence and quantify the samples, they were loaded into the MiSeq equipment.

A code-based approach (barcoding) was used for the complete analysis of the gut microbiome using the OTUs grouping methods. OTU is defined as organisms grouped by similarities in their DNA sequence, with a sequence similarity threshold of at least 75% to 80%. The taxonomy was assigned using BLAST and HITdb, and the sequences were filtered following the OTU LotuS quality criteria (version 1.58). The abundance matrices were filtered and then normalized at each level of classification: OTU, species, genus, family, order, class, and phylum.

In order to perform the comparative analyses to evaluate the differences between gut microbiota composition the MicrobiomeAnalyst tool (https://www.microbiomeanalyst.ca/) was used. For these analyses, the raw count of microorganisms was used. The DESeq2 (RNA-seq methods) analysis was used for finding significant differences in abundance of species and genera, using the trimmed mean of M-values (TMM) normalization and assigning the SILVA taxonomy labels. DESeq2 is a robust method that shows low false positive rates and according to recent guidelines, it has the highest power to compare groups, especially for less than 20 samples per group (Weiss et al., 2015).

### Humans CSF samples

Cerebrospinal fluid (CSF) samples were collected in a study conducted by the Karolinska University Hospital (Sweden). 22 patients with subjective cognitive impairment (SCI) were grouped as controls group because they didn’t present complaints on objective cognitive tasks. CSF samples were obtained by lumbar puncture from L3/L4 o L4/L5 interspaces in the mornings. After disposal of the first milliliter, the following 10 mL were collected in polypropylene tubes. No sample containing more than 500 erythrocytes/μL of CSF was used. Samples were gently mixed to avoid gradient effects and centrifuged at 2000 x g at 4°C for 10 min to eliminate cells an insoluble material. Supernatants were aliquoted, immediately frozen and stored at −80°C.

### TMAO measurement by UPLC-MS/MS

Serum and CSF TMAO levels were quantified using ultra performance liquid chromatography-tandem mass spectrometry (UPLC-MS / MS) with isotopic marking, using deuterated TMAO standard as described previously (Zhao, Zeisel, and Zhang 2015). Briefly, 20 μL of serum were added to a 1.5 ml Axygen tube containing 80 μL of internal standard mixture of 0.5 μM d9-TMAO in acetonitrile/methanol/formic acid (75/25/0.2 v/v). Once mixed, the sample was vortexed for 30 seconds to precipitate proteins following a centrifugation at 12000 rpm at 4 °C for 15 minutes to recover the supernatant. To elaborate the standard curve, nine samples of known concentrations of TMAO ranging from 0.15 μM to 60 μM were processed using the same procedure reaching a coefficient of determination (R_2_) above 0.999. Finally, 85 μL of the supernatant were diluted with 20 μL of MilliQ water and once mixed, it was analysed by injection onto a pre-column (X-Bridge BEH Amide 2,5µm, VanGuard Precolumn 2,1 × 5mm, Waters) following a column (X-Bridge BEH Amide 2,5 µm 2,1 × 50mm, Waters) at a flow rate of 0.4 mL/min using UPLC-MS/MS (Waters).

A discontinuous gradient was generated to resolve analytes using phase A (Amonium Acetate 10mmM in MilliQ Water) and phase B (Amonium Acetate 10mmM in acetonitrile/MilliQ water 90/10 v/v) as follows:

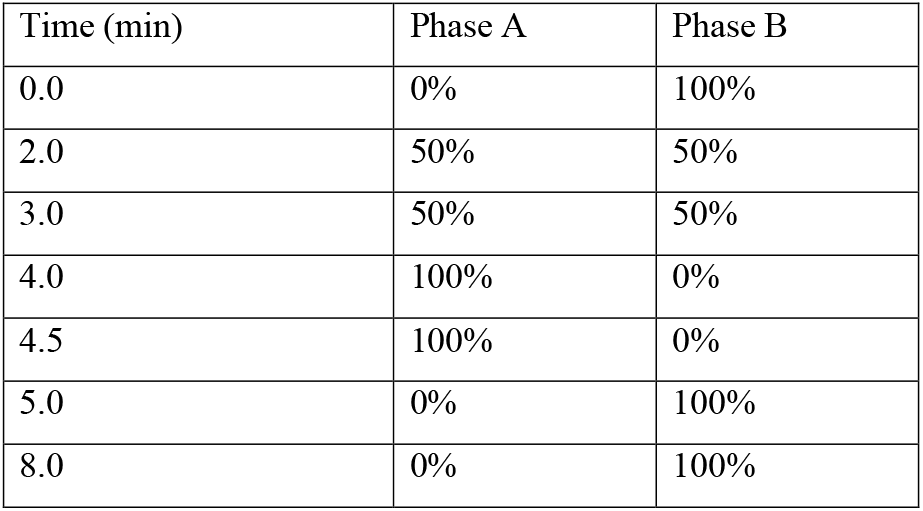

Analytes were monitored using electrospray ionization (ESI) in positive-ion mode [M-H]+ with multiple reaction monitoring (MRM) of precursors and characteristic production transitions of TMAO at m/z 76→58, d9-TMAO at m/z 85→66, respectively.

For the measurement of TMAO in brain tissue samples approximately 40 mg of parietal cortex were added to a 1.5 ml Axygen tube containing 120 μL of internal standard mixture of 0.5 μM d9-TMAO in MilliQ water. Samples were sonicated in ice with short pulses to homogenize. Supernatant was recovered and 50 μL were mixed with 100 μL of acetonitrile with 1% formic acid. After briefly mixing, the mixture was vortexed for 30 seconds to precipitate proteins following a centrifugation at 12000 rpm at 4 °C for 15 minutes to recover the supernatant. Finally, the supernatant was filtrated using OSTRO plates to get rid of phospholipids and proteins and injected in the column as previously described. To elaborate the standard curve, nine samples of known concentrations of TMAO ranging from 0.04 μM to 4 μM were processed using the same procedure reaching a coefficient of determination (R_2_) above 0.999.

### Statistical analysis

Results, reported as means ± SEM. Normality was checked by Shapiro–Wilk’s test (p<0.05). Statistical analysis of MWM acquisition, as well as, ITT and GTT tests was performed by repeated measures ANOVA. Rest of the data were analysed by two-way ANOVA tests followed by Tukey’s using GraphPad Prism, version 6. p<0.05 was considered to indicate statistical significance. Correlation test for CSF human samples was carried out by Spearman test.

## Results

### Aging alters peripheral insulin sensitivity and glucose homeostasis in SAMP8 mice but not brain insulin signaling

Peripheral insulin sensitivity and glucose homeostasis were assessed by ITT and GTT tests. SAMP8 mice showed enhanced insulin resistance (repeated measures ANOVA, main effect of strain, F_5,15_=5.536; *p*<0.01) (Figure 1a and b) and glucose intolerance (repeated measures ANOVA, main effect of strain, F_5,20_= 2.263; *P*<0.05) when compared to SAMR1 mice (Figure 1c and d). Moreover, blood glucose levels increased with age in both strains (two-way ANOVA, main effect of age, F_2,58_=26.21; *p*<0.0001) but in all time points SAMP8 mice showed elevated glucose concentrations (two-way ANOVA, main effect of age, F_1,58_=6.319; *p*<0.01) compared to SAMR1 mice (Figure 1e). Surprisingly, concerning insulin and HOMA index only a main effect of age was observed (Two-way ANOVA, F_2,58_=26.21; *p*<0.0001 and Two-way ANOVA, F_2,52_=0.1630; *p<*0.001, respectively), but although a strong tendency can be observed (especially in the HOMA index), no effect of strain was found, probably due to the high variability of the data (Figure 1f and g).

**Figure 1.**
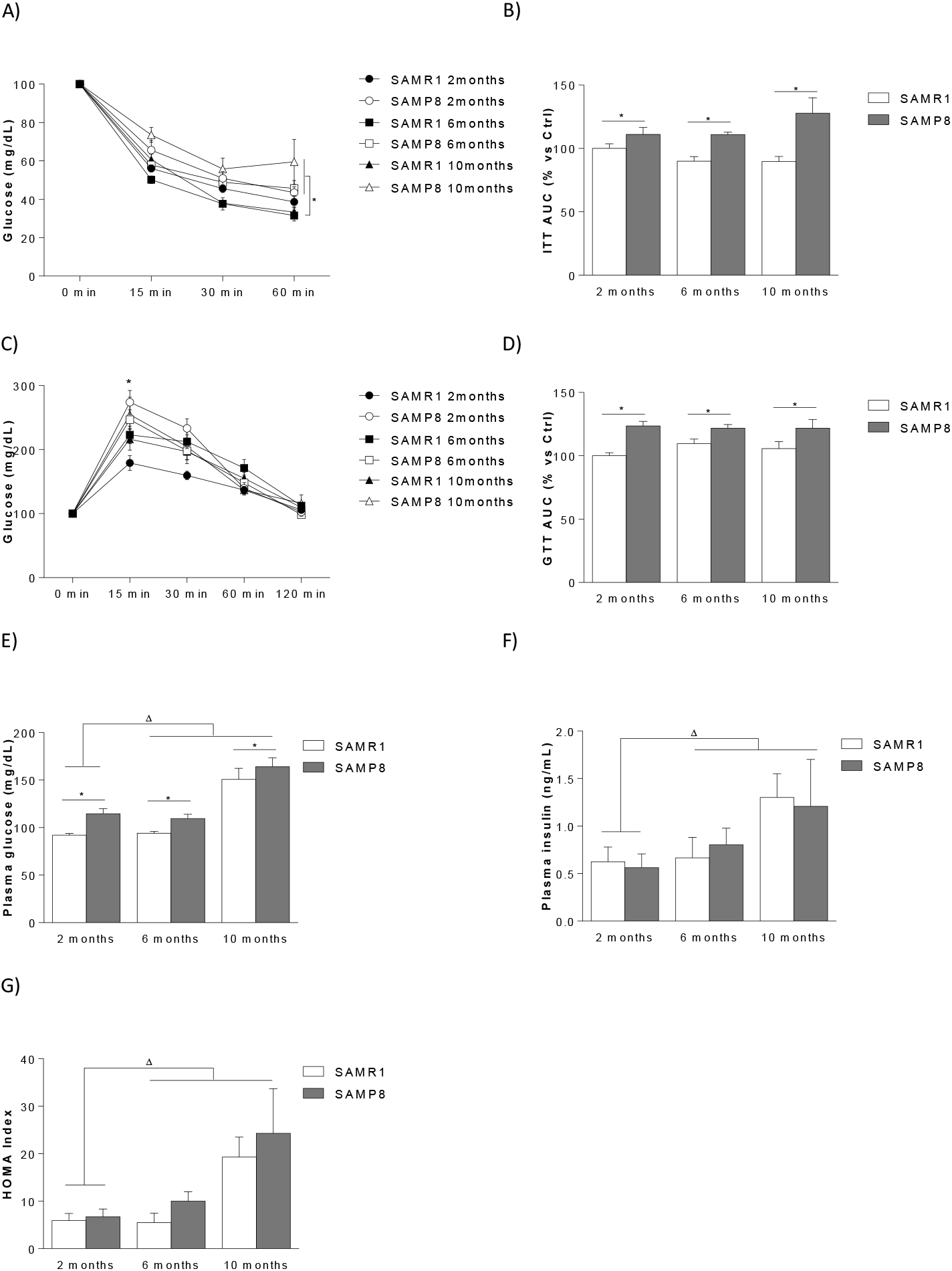
Effect of aging on peripheral insulin sensitivity and glucose homeostasis. The following metabolic parameters were analysed in all six groups. A) Insulin tolerance test (ITT). B) ITT expressed as area under curve (AUC). C) glucose tolerance tests (GTT). D) GTT expressed as area under curve. E) blood glucose levels F) blood insulin levels G) homeostatic model assessment indices of insulin resistance (HOMA). Data are presented as mean ± SEM. *Main effect of strain, two way ANOVA; ^Δ^ Main effect of age, two way ANOVA.

The observed peripheral metabolism alteration was not due to an increase in SAMP8 mice size or body weight. Indeed, SAMP8 body weight was found to be significantly lower when compared to same age SAMR1 littermates (two-way ANOVA, main effect of strain, F_1,61_=152.4; *p*<0.0001), although body weight is progressively increased in both strains with the age (two-way ANOVA, main effect of age, F_2,61_=57.66; *p*<0.0001) (Suppl. Figure 1a). In order to discover if differences in body weight are due to alterations in body composition, fat and lean mass content were measured in mice. No significant differences were observed between groups in lean (two-way ANOVA, F_2,61_=2.174; *p*=0.1225) neither in fat mass (two-way ANOVA, F_2,61_=2.837; *p*=0.0763) (Suppl. Figure 1b).

In order to study the possible insulin signalling alterations in the brain, expression of several components of the insulin signalling were analysed. Only 10 months SAMP8 mice group showed significant changes in pIRS (Tukey’s p<0.05 vs rest of the groups) (Suppl. Figure 2b) and pAkt levels (Tukey’s p<0.05 vs rest of the groups) (Suppl. Figure 2c). However, no changes were found in the rest of the protein studied (pIR, Suppl. Figure 2a; pGSK3β, Suppl. Figure 2d; pERK1, Suppl. Figure 2e; pERK2, Suppl. Figure 2f), suggesting that there is not brain insulin resistance. Consistent with a post-transcriptional regulation of these enzymes, total protein levels, normalized using actin, remained unaltered.

### Aging induces cognitive deficiencies in SAMP8 mice

As it was expected, age dependent locomotor activity decrease was found (two-way ANOVA, main effect of age, F_2,58_=15.45; *p*<0.0001) (Figure 2a). Interestingly, there were no differences between strains, indicating that behavioural performance differences between SAMP8 and SAMR1 are not due to locomotor activity alterations.

**Figure 2.**
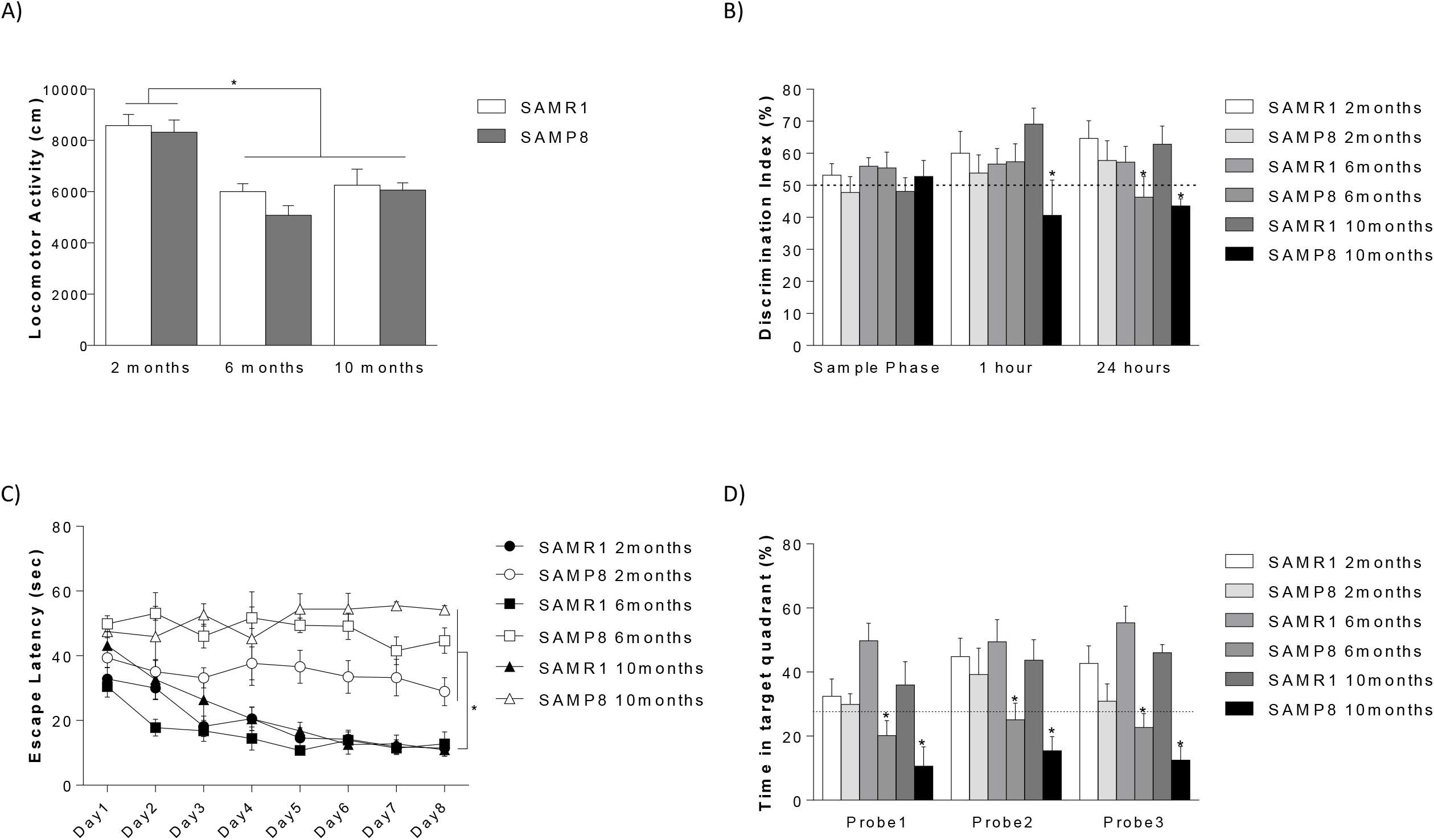
Effect of aging on cognitive performance. In panel A) locomotor activity. In panel B) cognitive performance in novel object recognition test (NORT). Data shows discrimination index (time exploring the new object / total exploration time × 100) in the novel object recognition test. In panel C) and D) cognitive performance assessed by Morris water maze (MWM) acquisition phase and retention phase respectively. Data are presented as mean ± SEM. *Tukey’s multiple comparison test, p<0.05.

As shown in figure 2b, a main effect of the strain was found (two way ANOVA, main effect of strain, F_5,163_=2.635; *p*<0.05). Further analysis showed that SAMP8 10 month mice displayed cognitive deficits in the NORT, as shown by a significantly decreased discrimination index in the 1 and 24 hour task (Tukey’s *p*< 0.05 vs rest of the groups). Cognitive impairment was also observed in SAMP8 6 months mice in the 24 hour NORT task (Tukey’s *p*<0.05).

In the acquisition phase of Morris water maze, distance swam to reach the platform improved significantly over trials in SAMR1 group (repeated measures ANOVA, F_2,14_=5.596; *p*<0.05) but not in SAMP8 group (repeated measures ANOVA, *p*>0.05). Moreover, significant effect of strain was found (repeated measures ANOVA, main effect of strain, F_5,35_=52.40; *p*<0.0001) as SAMP8 mice in all ages showed higher scape latency compared to SAMR1 mice indicating a cognitive impairment (Figure 2c). In the retention phase, 6 and 10 months SAMP8 mice showed a statistically significant decrease in time swam in the quadrant were the platform used to be located, indicative of a memory deficit (Tukey’s *p*<0.05 in all cases) (Figure 2d).

### Aging disrupts BBB integrity in SAMP8 mice

BBB integrity was evaluated by immunofluorescence staining of fibrin and IgG in the left hemispheres of SAMR1 and SAMP8 groups. Fibrin (two way ANOVA, main effect of strain, F_1,18_=5.730; *p*<0.05) (Figure 3) and IgG (two way ANOVA, main effect of strain, F_1,12_=4.641; *p*<0.05) (Suppl. Figure 3) extravasation was increased in SAMP8 6 and 10 months aging groups compared to SAMR1 mice.

**Figure 3.**
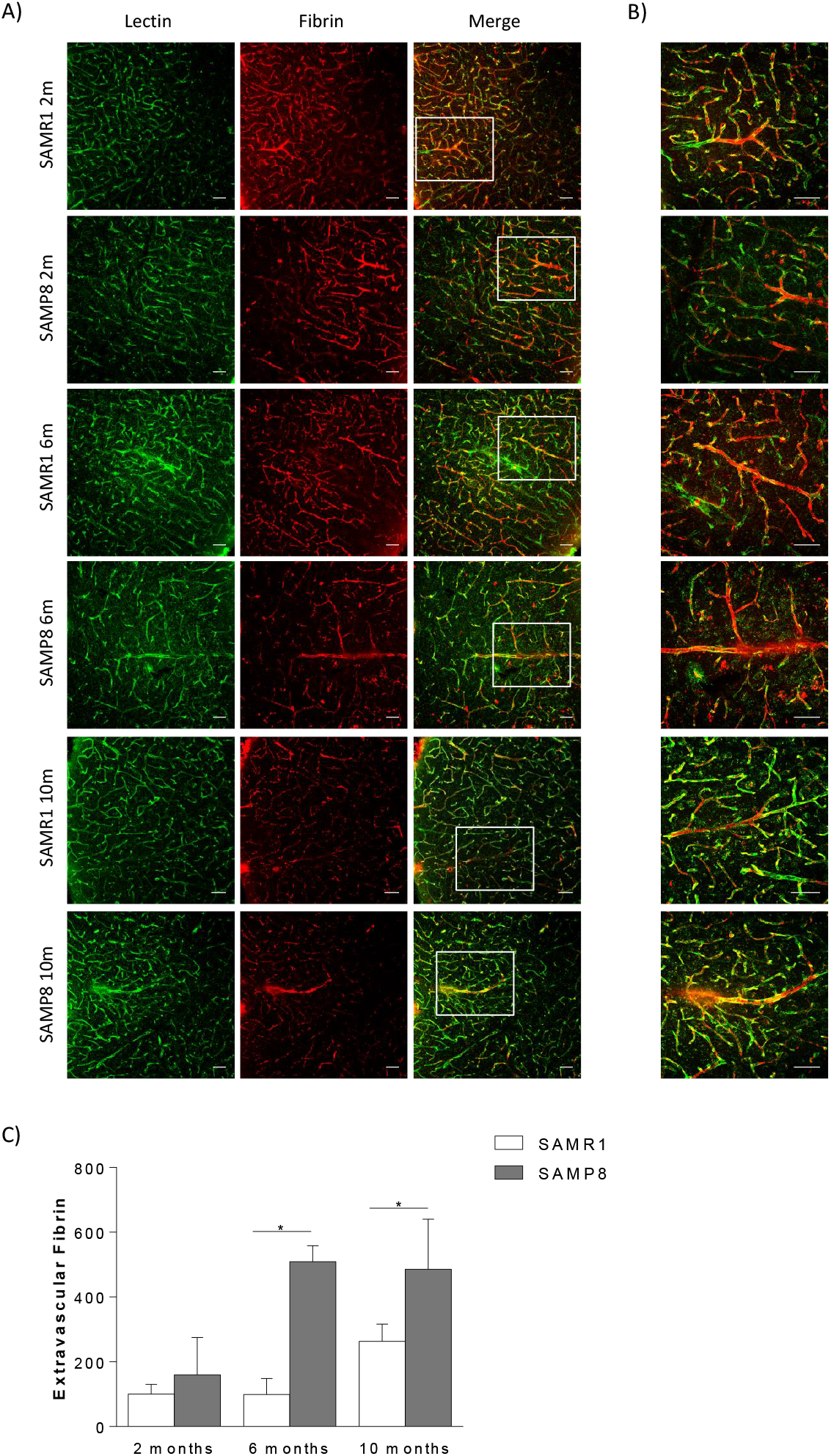
Effect of aging on BBB integrity. Representative confocal microscopy analysis of A) fibrin (red) and lectin-positive capillaries (green) on cortical brain sections and B) amplification of selected areas. In panel C) quantification of the extravascular fibrin. * Main effect of strain, two way ANOVA, p<0.05. Scale bar, 100 μm.

Moreover, protein levels of two of the main components of the tight junctions (occludin and ZO-1) appeared to be significantly decreased in SAMP8 mice compared to SAMR1 mice (for occludin, two way ANOVA, main effect of strain, F_1,30_=11.31; *p*<0.01; for ZO-1, two way ANOVA, main effect of strain, F_1,30_=20.46; *p*<0.0001) (Figure 4).

**Figure 4.**
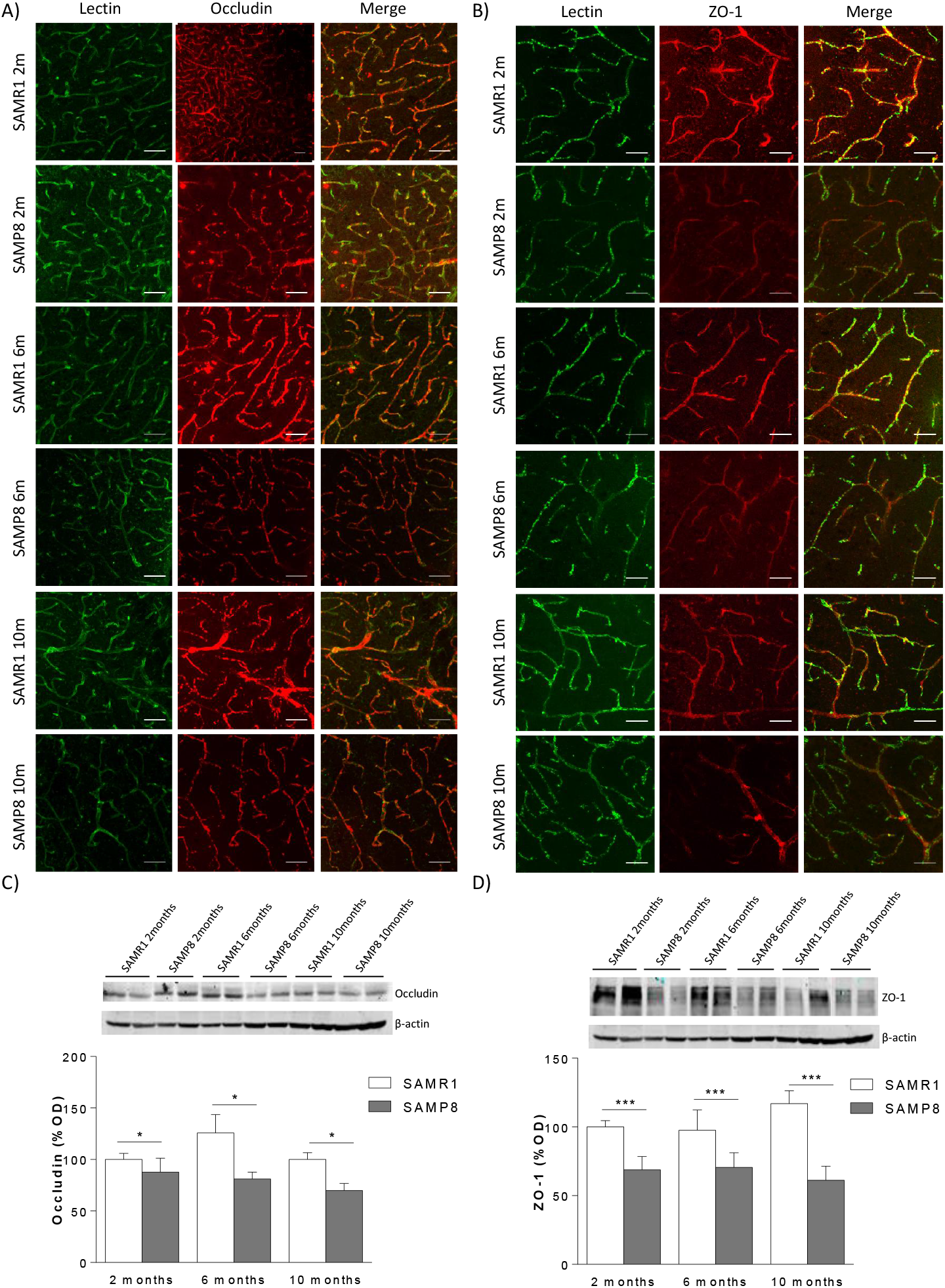
Effect of aging on BBB tight junctions. Representative confocal microscopy analysis of A) occludin (red) and B) ZO-1 (red) and lectin-positive capillaries (green) on cortical brain sections. In panel C) Occludin and D) ZO-1 representative immunoblots from mice brain tissue and their quantification. * Main effect of strain, two way ANOVA, p<0.05. Scale bar, 100 μm.

### Aging promotes peripheral and central inflammation

Our data showed an increase in inflammatory cytokines (TNFα and IL6) not only in the periphery (white adipose tissue) (for TNFα, two way ANOVA, main effect of strain, F_1,18_=6.534; *p*<0.05; for IL-6, two way ANOVA, main effect of strain, F_1,18_=6.639; *p*<0.05) (Figure 5a and b) but also in the central nervous system (Frontal cortex, Figure 5c and d) of SAMP8 mice (for TNFα, two way ANOVA, main effect of strain, F_1,18_=4.570; *p*<0.05 and main effect of age, F_1,18_=4.708; *p*<0.05; for IL-6, two way ANOVA, main effect of strain, F_1,18_=8.367; *p*<0.01). Neuroinflammation was further confirmed by pJNK and GFAP (an astrocyte activation marker) protein levels. As depicted in Suppl. Figure 4a, a significant increase in pJNK levels in SAMP8 mice in all ages were observed (two-way ANOVA, main effect of strain, F_1,42_= 27.82; *p*<0.0001). In addition, an age and strain dependent significant increase in GFAP levels were observed in SAMP8 mice compared to SAMR1 group (two-way ANOVA, main effect of age, F_2,41_=14.05; *p*< 0.0001; two-way ANOVA, main effect of strain, F_1,41_=6.327; *p<* 0.05) (Suppl. Figure 4b). The increase in GFAP was also observed by immunohistochemistry analysis (Suppl. Figure 4c).

**Figure 5.**
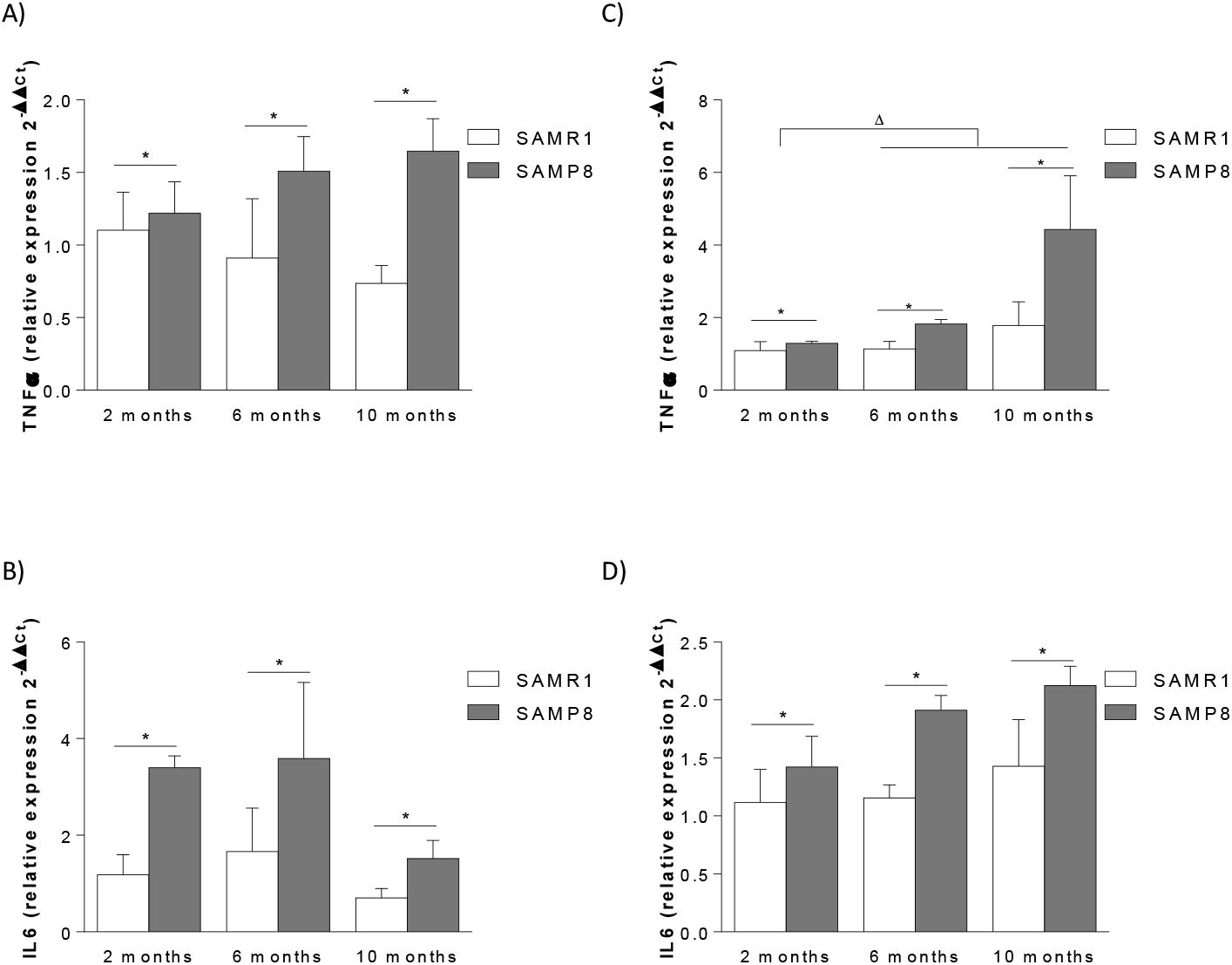
Effect of aging on pro-inflammatory cytokine release. Peripheral (white adipose tissue) A) TNFα and B) IL6 gene expression. Cortical (frontal cortex) C) TNFα and D) IL6 gene expression. *Main effect of strain, two way ANOVA; ^Δ^ Main effect of age, two way ANOVA.

### Aging induces gut microbiota dysbiosis

α-diversity is a measure of microbiome diversity within a community and is mainly concerned with the number of different bacteria or species therein. Shannon and Simpson are indices commonly used to evaluate α-diversity of microbiota. Shannon (p=0.012904, Suppl. Figure 5A) and Simpson (p=0.041409, Suppl. Figure 5B) indices were significantly lower in fecal samples from SAMP8 mice, thus indicating less diversity of bacteria. On the other hand, β-diversity is an estimate of similarity or dissimilarity between populations. PCoa-Jensen Shannon Divergence analysis revealed that dots from SAMR1 mice were not close to those of SAMP8 (R=0.333, p<0.05), suggesting that SAMP8 have a different microbiome composition than SAMR1 (Suppl. Figure 5C). Specifically, SAMP8 mice showed a significant increase in Bacteroidetes phylum (Figure 6A) that was probably due to an increase in Prevotella genus (Figure 6B). Moreover, a profound decrease in Proteobacteria phylum was found (Figure 6C), with a marked decrease in Deltaproteobacteria class (Figure 6D). Although no changes were observed in the Firmicutes phylum (Figure 6E), it is worth mentioning a marked decrease in Dorea genus (Figure 6F). Finally, although the Actinobacteria phylum appeared unchanged in SAMP8 mice (Figure 6G), Actinobacteria class showed a significant increase (Figure 6F).

**Figure 6.**
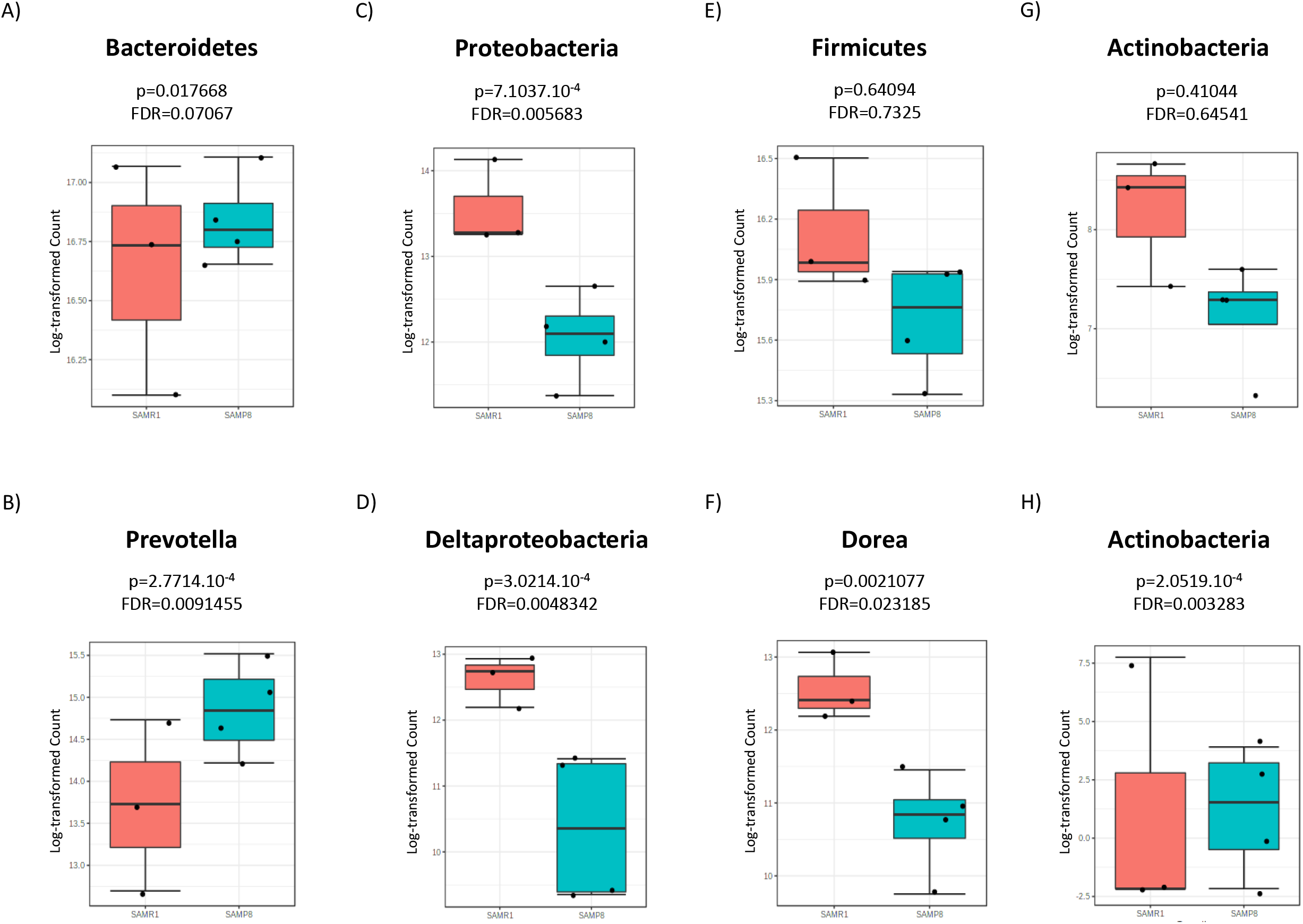
Effect of aging on gut dysbiosis. Bacterial phylum, class and genera that are significantly changed (edgeR p value <0.05 and FDR <0.05) between SAMPR1 and SAMP8 mice groups. Data were log-transformed counts of bacterial 16 S rRNA gene copies.

### Aging increases peripheral and central TMAO levels

As depicted in figure 7A, aging induces a significant increase in plasma TMAO levels of SAMP8 mice (two way ANOVA, F_2,30_=9.878; p<0.001 followed by Tukey’s p<0.05 SAMP8 vs rest of the groups). Interestingly, elevated levels of TMAO are also observed in the aging brain tissue of SAMP8 mice (two way ANOVA, main effect of age F_2,31_=62.43; p<0.0001) (Figure 7B). In the same line, human CSF TMAO levels showed a positive correlation with aging (Spearman r=0.5793, p<0.01) (Figure 7C) and BBB integrity measured by brain albumin quantity (Spearman r=0.6740, p<0.01) (Figure 23D).

**Figure 7.**
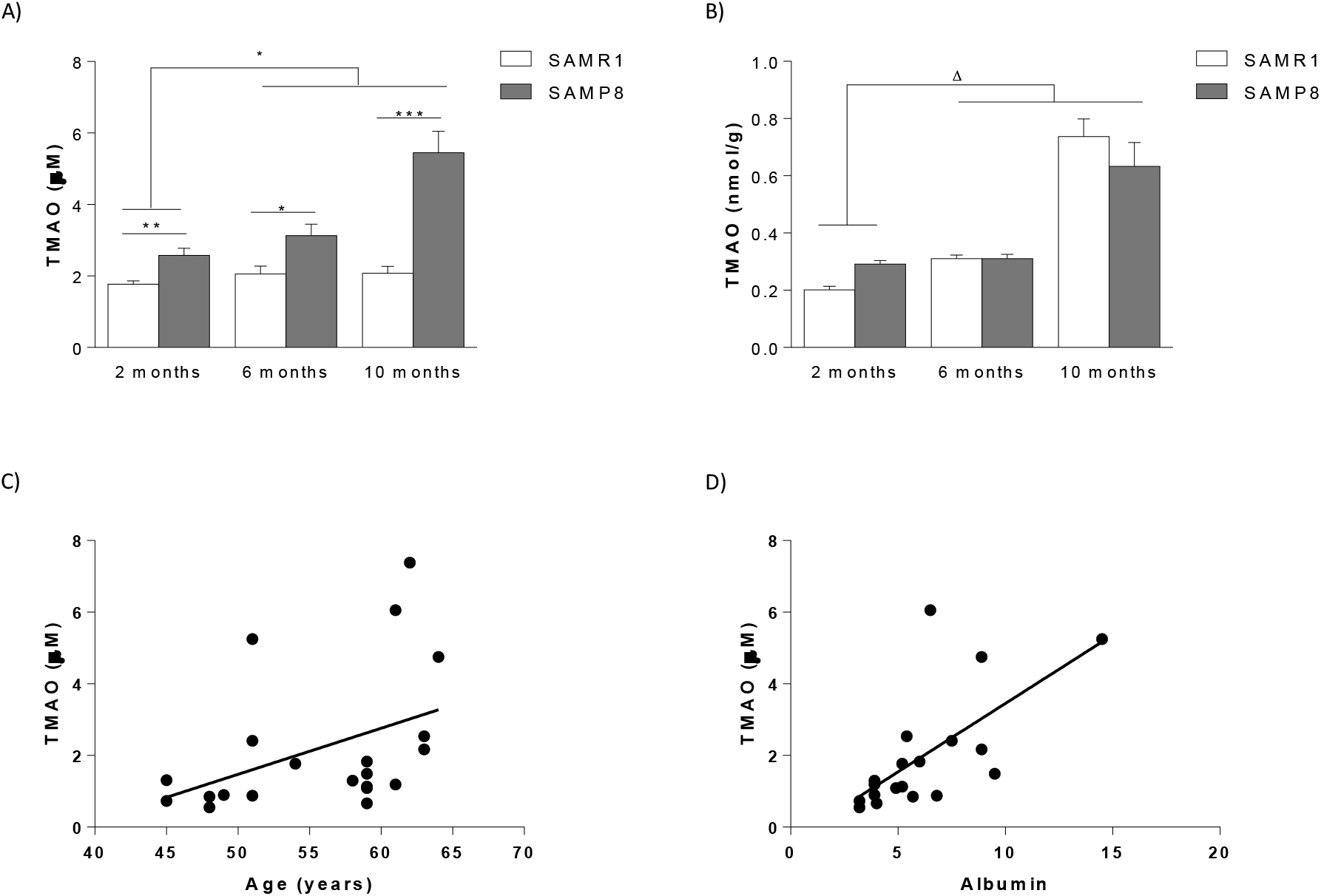
Effect of aging on TMAO levels. In panel A) TMAO levels in plasma samples in mice and B) TMAO levels in mouse brain samples. Human CSF TMAO levels correlate with age (panel C) and BBB integrity (panel D).

### DMB treatment restores TMAO-induced alterations in SAMP8 mice

Treatment with DMB, choline TMA lyase enzyme inhibitor that decreases TMAO plasma levels, showed clear improvement of peripheral metabolism, as SAMP8 mice treated with DMB showed reduced adiposity (two way ANOVA, main effect of treatment F_1,38_=6.449; *p*<0.05) (Figure 8A), improved glucose tolerance (repeated measures ANOVA, F_12,124_= 2.567; *p*<0.01) (Figure 8B), lower fasting plasma glucose (two way ANOVA, main effect of treatment F_1,26_=12.99; *p*<0.01) (Figure 8C) and insulin levels (two way ANOVA, main effect of treatment F_1,18_=7.444; *p*<0.01) (Figure 8D), that leads to a decreased HOMA index (two way ANOVA, main effect of treatment F_1,18_=37.13; *p*<0.0001) (Figure 8E). In parallel, DMB treated SAMP8 mice showed a significantly improved performance in the acquisition Morris water maze test (repeated measures ANOVA, F_3,29_= 24.80; *p*<0.0001) (Figure 8F) as well as in the second probe trial (two way ANOVA, F_1,23_=6.644; *p*<0.05, followed by Tukey, p<0.05 SAMP8 vehicle vs SAMR1 vehicle) (Figure 8G). This cognitive enhancement was accompanied by a decrease in brain neuroinflammatory markers, i.e., lower TNFα (two way ANOVA followed by Tukey’s p<0.05 SAMP8 vs SAMP8 DMB) (Figure 8H), p-JNK (two way ANOVA, main effect of treatment F_1,24_=6.462; *p*<0.05) (Figure 8I) and GFAP levels (two way ANOVA followed by Tukey’s p<0.05 SAMP8 vs SAMR1 vehicle) (Figure 8J).

**Figure 8.**
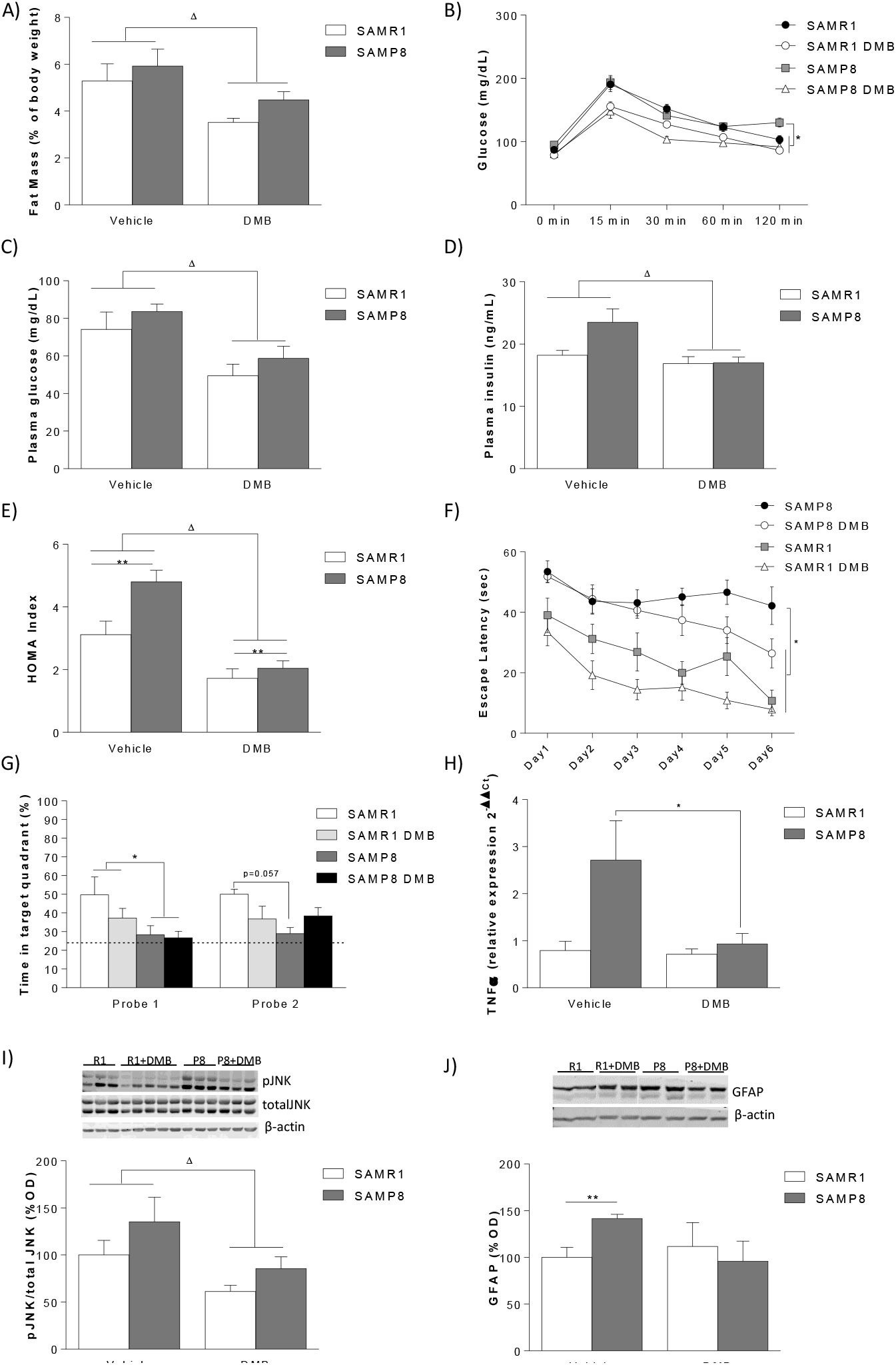
Effect of DMB on SAMP8 mice. In panel A) mice adiposity measurement. B) Glucose tolerance tests (GTT) after treatment with DBM. C) Serum glucose, D) serum insulin and E) Homeostatic model assessment indices of IR (HOMA) after treatment with DBM. In panel Cognitive performance assessed by Morris water maze (MWM) F) acquisition phaes and G) retention phase. Cortical (frontal cortex) H) TNFα expression, I) p-JNK protein levels and J) GFAP protein levels.. *Main effect of strain, two way ANOVA; ^Δ^ Main effect of treatment, two way ANOVA.

## Discussion

In recent decades, significant advances in modern medicine, coupled with an improved quality of life in developing countries, have significantly increased life expectancy and older people are expected to live even longer in the future (WHO, 1998). The improvement in life expectancy is inevitably accompanied by a subsequent escalation in the prevalence of age-related diseases. Among them, insulin resistance and neurodegenerative diseases such as Alzheimer’s disease (AD) are considered part of the main threats to health in old age (Ballard et al., 2011; Mokdad et al., 2001).

Classically, insulin resistance and neurodegeneration have been considered unrelated pathological entities, either as a metabolic disorder that mainly affects glucose homeostasis in peripheral organs such as the skeletal muscle, liver and fat, or as a degenerative disease of the central nervous system (CNS), respectively. However, recent studies have raised the possibility that these diseases may have similar molecular roots. Indeed, recently, both diseases have been associated with impaired action of insulin in the CNS. This notion is supported by epidemiological studies that have found a link between insulin resistance and AD (Frisardi et al., 2010). However, other studies have not been able to reveal this relationship (Frisardi et al., 2010). Post-mortem analyzes of the brains of AD patients have shown that insulin receptors are downregulated (Frolich et al., 1998), as observed during aging (Fernandes et al., 2001; De Felice, 2013). This led to the hypothesis that neuronal insulin resistance may contribute to the etiology of AD. Therefore, it is believed that the close correlation between metabolic disturbances (such as diabetes mellitus) and cognitive deficits is mainly due to the establishment of insulin resistance and the associated changes in the pleiotropic effects of insulin on central physiological functions. Therefore, the aim of the present project was to investigate as the missing link between aging, insulin resistance and dementia in a mouse model of accelerated senescence (SAMP8) in an age fashion (2, 6 and 10 months).

Aging is associated with marked alterations in insulin secretion, which commonly leads to hyperinsulinemia (Fernandes et al., 2001). In our hands, marked insulin impairment and glucose intolerance that worsen with the age was found in SAMP8 mice group. This together with a strong tendency of increased HOMA index indicates the existence of age related peripheral insulin resistance and altered glucose homeostasis. The critical involvement of abnormal insulin signaling in the etiology of metabolic diseases has been emphasized by the onset of mild obesity and altered glucose metabolism in neuronal insulin receptor knock out (NIRKO) mice (Bruning et al., 2000). Although the exact molecular mechanisms driving the onset of insulin resistance are not yet fully understood, it is evident that two of the main risk factors for development of insulin resistance are overweight/obesity and aging. Therefore, next step was to analyze if aging per se induces body weight changes and fat accumulation that could lead to peripheral insulin resistance. Body weight was increased equally over time in both strains studied in the present work but it was always lower in SAMP8 mice compared to SAMR1 due to their smaller size. Surprisingly, as aging induces fat content accumulation a higher fat mass was expected in SAMP8 mice; however, no differences in body composition were found between groups. Therefore, the observed insulin resistance is not related to a higher adiposity, which suggests that an alteration in the central nervous system driven insulin signaling regulation could be responsible of this defect.

Interestingly, in parallel with the observed peripheral insulin resistance, SAMP8 mice showed a marked age dependent cognitive decline, at least in the Morris water maze, a hippocampal-dependent task. The results found on the NORT, where only 10 months old SAMP8 mice exhibited cognitive impairment in the 1 hour task together with the fact that the hippocampus is only minimally involved in memory for objects (Brown and Aggleton, 2001), would support the notion that memory deficits in aging firstly has its anatomical substrate on the hippocampus and only when elderly is reached cortex starts been affected. It is worth mentioning that even though 6 and 10 months mice seem to have a decreased activity in the open field test, other measurements of locomotor activity, such as swim speed in the Morris water maze or total exploration time in the first exposure to objects in the NORT, did not differ among groups.

Almost a century after the discovery of insulin as a peptide secreted by the pancreas (Banting and Best, 1922), the classical view of insulin as one of the main regulators of glucose homeostasis by promoting glucose uptake in peripheral tissues, such as the muscle and fat pads, and suppressing hepatic glucose production has expanded to a wide range of physiological and cellular effects, including its neuroprotective function and the regulation of learning and memory (Fernandez and Torres-Aleman, 2012). Based on this idea, we next studied if changes in insulin signalling in SAMP8 mice observed in the periphery were also occurring in the CNS and could be responsible of the cognitive deficits. Previous studies have shown that insulin sensitivity in the CNS is regulated in an age dependent manner. As we age, many of the steps that control insulin action change, from alterations in the level of insulin itself to its intracellular signaling pathways. In fact, in addition to the gradual generalized loss of basic physiological functions with age, the decrease in insulin action appears to be an inevitable consequence of aging (Kushner et al., 2013). The expression of insulin receptors in the brain is also subjected to an age-dependent decline (Zhao et al., 2004). Notably, the level of insulin receptor mRNA in the hypothalamus, cortex, and hippocampus of old rats is drastically reduced (Zhao et al., 2004). Surprisingly, our data showed a slight decrease in pIRS1 and pAkt levels in 10 months SAMP8 mice without alteration in any other component of the brain insulin signalling. Thus, we can concluded that the peripheral insulin resistance observed in SAMP8 mice is not transferred to the CNS and therefore the cognitive decline seen in those mice may not be due to central insulin resistance.

Given that central insulin resistance does not seem to be the cause of the observed cognitive disturbances, one can hypothesize the involvement of another molecular mechanism that, on one hand, leads to peripheral insulin resistance and, on the other hand, contributes to cognitive deficiencies, thus explaining the clinical link between peripheral metabolic disturbances and neurodegenerative disorders. In an attempt to find that missing mechanism underlying the age related cognitive deficiency observed in SAMP8 mice in the present study, we investigated other mechanisms that could be related to insulin signalling and could be the cause of a cognitive alteration. Mounting evidence has shown that the existing association between insulin signaling and the low-grade inflammatory tone observed in obesity may be one of the leading processes toward the onset of neuronal insulin resistance (Vogt and Bruning, 2013; Thaler et al., 2013; Velloso and Schwartz 2011; Konner and Bruning, 2011). Inflammation has been associated mainly with the realease of cytokines, particularly TNF-α and IL6. TNF-α activates inflammatory kinases such as c-Jun N-terminal kinase (JNK) (De Souza et al., 2005; Posey et al., 2009; Zhang et al., 2008). In addition to the detrimental effect of low-grade inflammation per se on cellular physiology, activation of inflammatory cascades blunt insulin receptor signaling directly, notably by interfering with the phosphorylation events downstream of insulin receptor activation (Vogt and Bruning, 2013). Specifically, activation of JNK leads to inhibition of IRS phosphorylation, and therefore to desensitization of insulin action (Vogt and Bruning, 2013) and (Aguirre et al., 2000). Interestingly, these aforementioned inflammatory pathways are also enhanced during normal aging (Zhang et al., 2013; Lee et al., 2000). In our hands, a marked peripheral inflammatory state (i.e. significantly higher adipose tissue TNF-α and IL6 levels) is observed in SAMP8 mice. The peripheral inflammation is accompanied by a strong neuroinflammation as SAMP8 mice exhibited significant increases of cortical and hippocampal TNF-α and IL6 levels together with an elevated pJNK levels as well as GFAP protein and inmmunoreactivity, already at 2 months of age. These results fits with the peripheral metabolism alterations and the cognitive deficiency observed in SAMP8 mice, suggesting that inflammation could be underlying both pathologies. Noteworthy, BBB integrity can be disrupted by neuroinflammation associated with many age-related disorders as Alzheimer’s disease (Cai et al., 2018). Indeed, SAMP8 mice showed elevated levels of brain fibrin and IgG accompanied by lower tight junction proteins expression, indicating an exacerbated BBB leaking. Thus, the aberrant amounts of peripheral pro-inflammatory cytokines observed may enter the brain via the BBB and cause inflammation by altering glial activation (Erny et al. 2015; Rothhammer et al. 2016) and subsequent brain cytokine reléase, leading to cognitive deficiencies.

Aging is a known risk factor for dysbiosis as it can alter gut microbiome composition and reduce microbiota diversity (Janeiro, Ramírez, and Solas 2021; O’Toole and Jeffery 2015). Dysbiosis can lead to a major production of certain gut metabolites such as TMA, which proceeds from the bacterial synthesis from substrates like choline or L-carnitine. It is then rapidly absorbed and further oxidized by hepatic enzymes (FMO1 and FMO3) to form TMAO (Janeiro et al. 2018). In our hands, a profound decrease in Proteobacteria, with a marked increase of Bacteroidetes was observed in SAMP8 mice. Bacteroidetes and specifically, *Prevotella* genus that appear to be significantly increased in SAMP8 mice have been associated with higher TMAO production. Indeed, literature has shown that individuals with an enterotype characterized by enriched proportions of *Prevotella* have significantly higher plasma TMAO, than individuals with a Bacteroides enterotype, indicating that enterotypes affect the host (Koeth et al. 2013). The higher TMAO levels could be also related to the observed decrease in the genus *Dorea*, as it has been previously reported in human samples where proportions of *Dorea* within recipient feces were inversely correlated with both plasma TMAO levels and atherosclerosis extent (Gregory et al. 2015). Furthermore, in parallel to our data, TMAO has been extensively positively correlated with the genus Bifidobacterium (phylum *Actinobacteria*) (O’Connor et al. 2014; Janeiro et al., 2018), concluding that the observed increases in this class could also lead to higher TMAO production.

In the last few years, TMAO has been widely associated with inflammation, glucose impairments and even neuropathologies (Janeiro et al. 2018). According to other studies (Ke et al. 2018), in the present study, an age-dependent increase of TMAO levels were observed not only in the periphery but also in the CNS, probably directly linked to the higher BBB leaking observed in SAMP8 mice. Therefore, it is temping to speculate that aging-induced dysbiosis can lead to higher TMAO levels inducing a systemic inflammatory status that may progress, in the context of a disrupted BBB, towards neuroinflammation, producing anomalous glial activation higher cytokine release and cognitive deficiencies in last instance (Janeiro, Ramírez, and Solas 2021).

In this context, a pharmacological treatment with 3,3-dimethyl-1-butanol (DMB) was established to reduce TMAO levels and ameliorate the cognitive decline seen at 6 months, prior to the huge increase of serum TMAO levels in SAMP8 mice. DMB is an analogue of choline that is able to reduce most but not all TMA lyases. It inhibits the transformation of choline, carnitine and crotonobetaine into TMA although it is not able to avoid the conversion of γ-butyrobetaine (GBB) (Wang et al. 2015). DMB treatment was able to reverse peripheral glucose alterations and to decrease HOMA index in SAMP8 mice. In parallel, cognitive performance was significantly improved and neuroinflammation appeared to be ameliorated.

Taken together, we can conclude that aging induced insulin resistance and dysbiosis with the subsequent elevated TMAO levels, can lead to an inflammatory state that could cause neuroinflammation on the one hand, and contribute to cognitive deficiency and AD like neurodegenerative progression on the other.

## Figure Legends

**Supplementary Figure 1.**
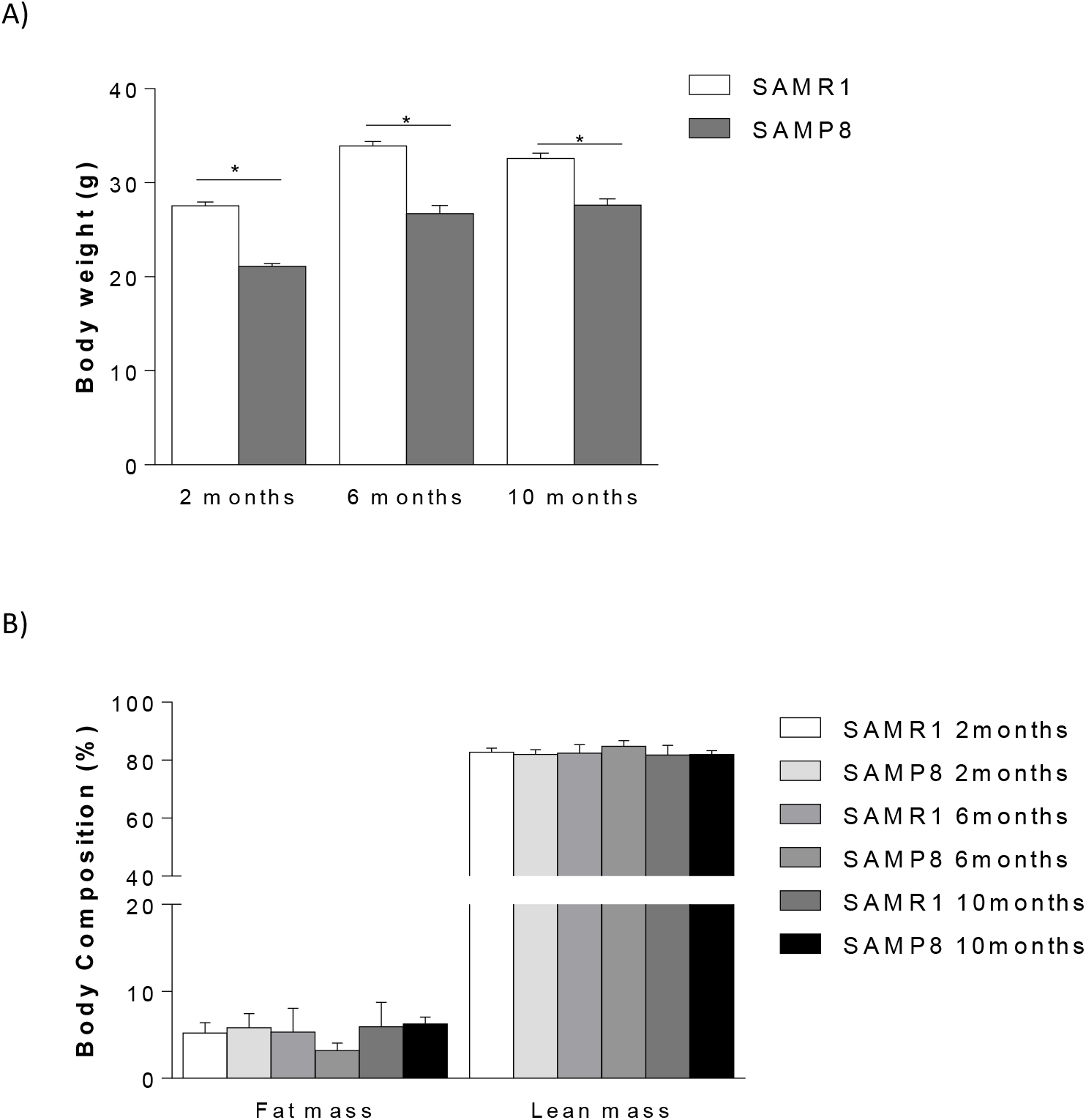
Effect of aging on body composition. In panel A) body weight. In panel B) mice body composition measurement with lean and fat mass assessment respectively. Data are presented as mean ± SEM. *Main effect of strain, two way ANOVA.

**Supplementary Figure 2.**
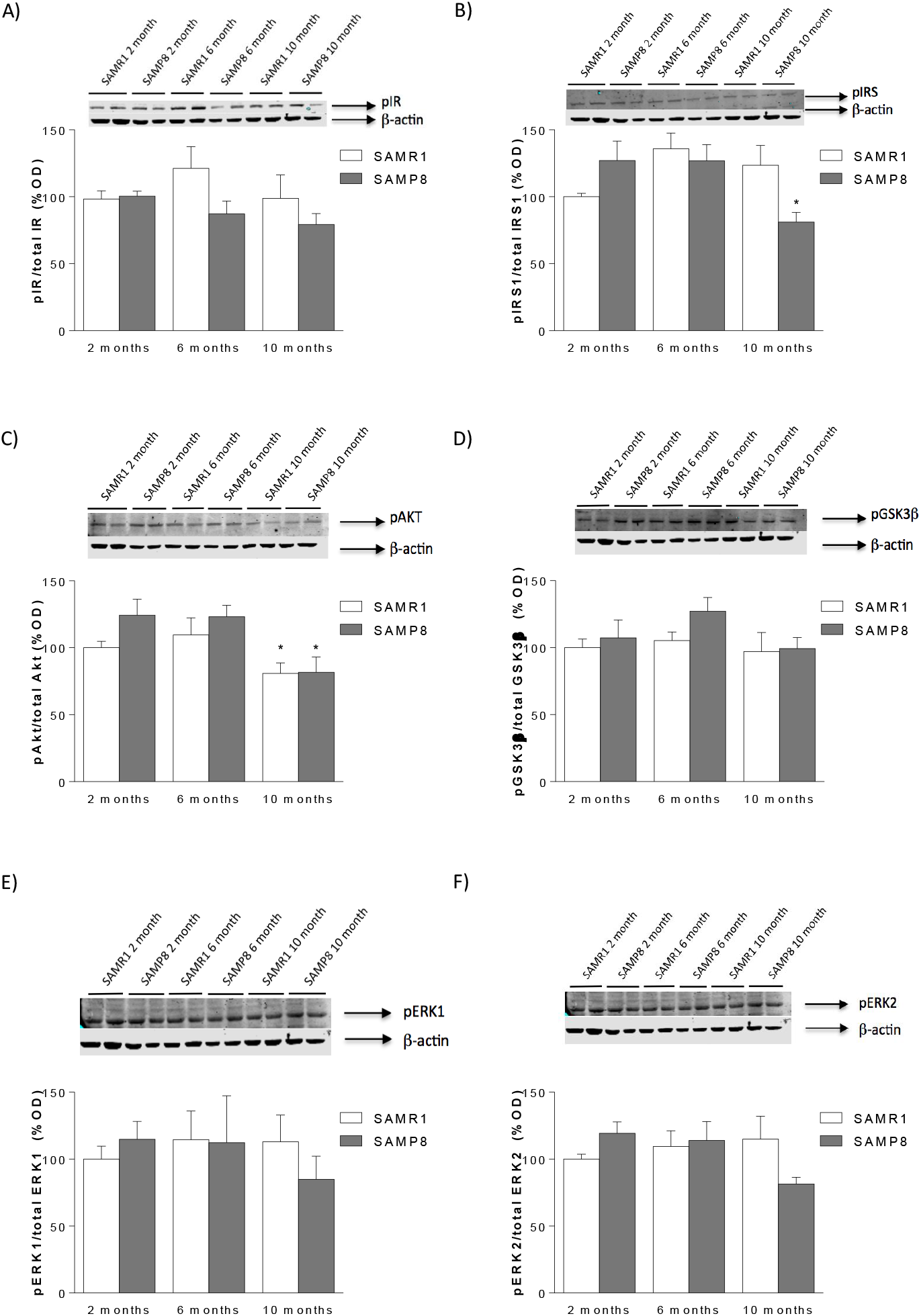
Effect of aging on insulin signalling in the central nervous system. Panel A) pIR expression levels. Panel B) pIRS1 expression levels. Panel C) pAKT expression levels. Panel D) pGSK3β expression levels. Panel E) pERK1 expression levels. Panel F) pERK2 expression levels. Figures show percentage of optical density (O.D.) values of control 2 months SAMR1 mice and representative picture of the blotting. No differences were found in the non-phosphorylated (total) levels of the enzymes. pIR: phosphorylated insulin receptor. *Tukey’s multiple comparison test, p<0.05.

**Supplementary Figure 3.**
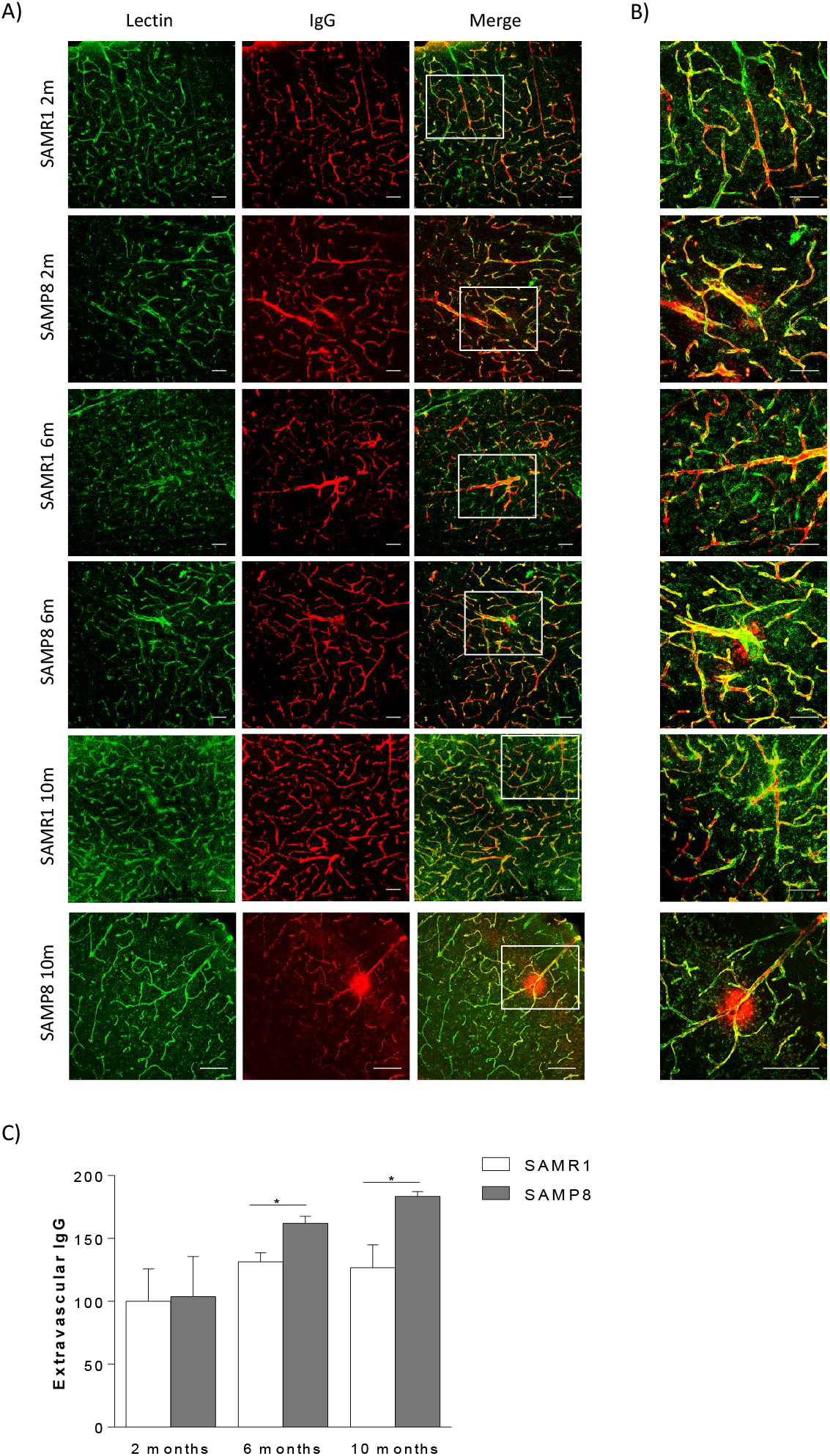
Effect of aging on BBB integrity. Representative confocal microscopy analysis of A) IgG (red) and lectin-positive capillaries (green) on cortical brain sections and B) amplification of selected areas. In panel C) quantification of the extravascular fibrin. *Main effect of strain, two way ANOVA, p<0.05. Scale bar, 100 μm.

**Supplementary Figure 4.**
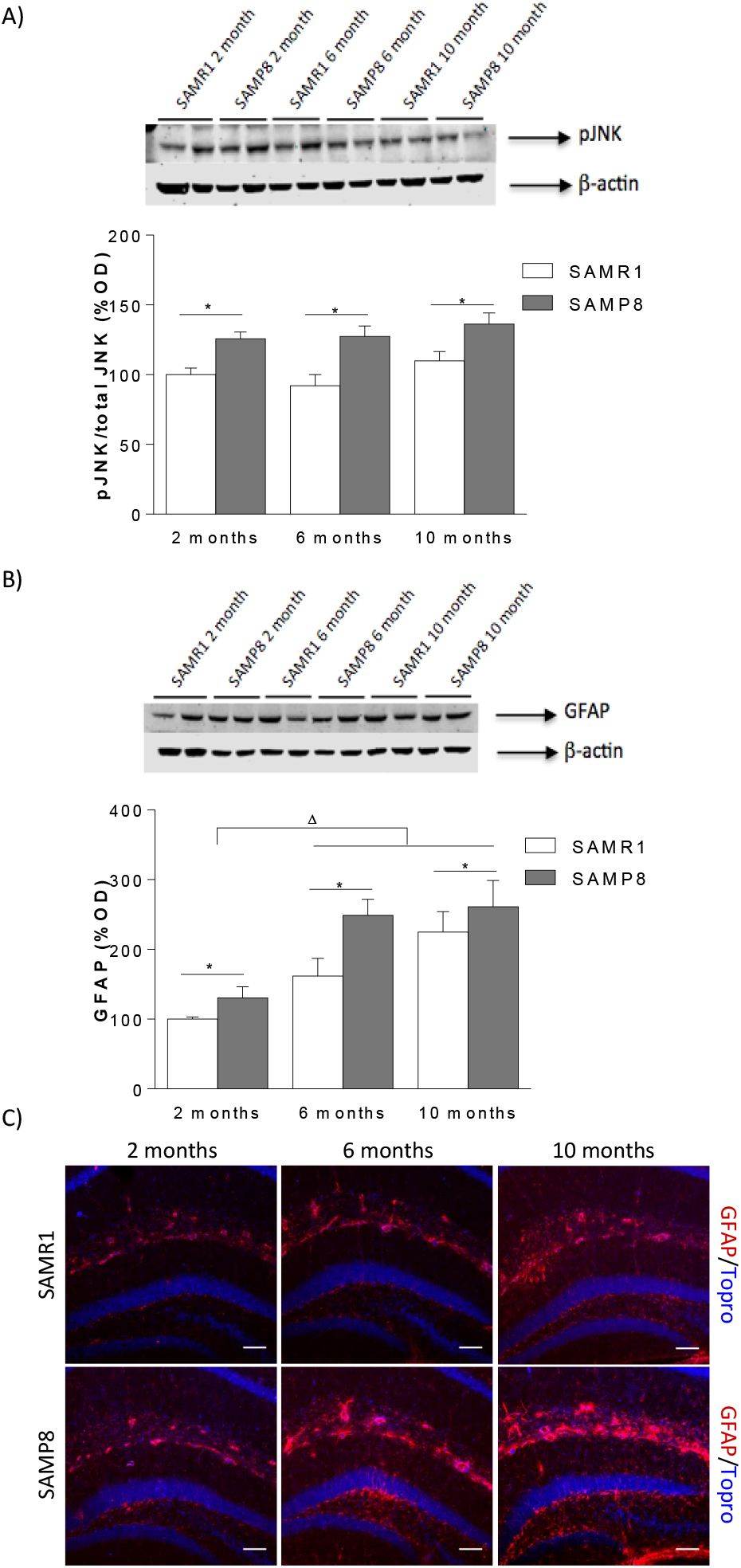
Effect of aging on neuroinflammation. Panel A) pJNK protein expression level. Panel B) GFAP protein expression levels. Panel C) GFAP immunohistochemical representative images. Scale bar 100 μm. Figures A and B show percentage of optical density (O.D.) values of control 2 months SAMR1 mice and representative picture of the blotting. *Main effect of strain, two way ANOVA; ^Δ^ Main effect of treatment, two way ANOVA.

**Supplementary Figure 5.**
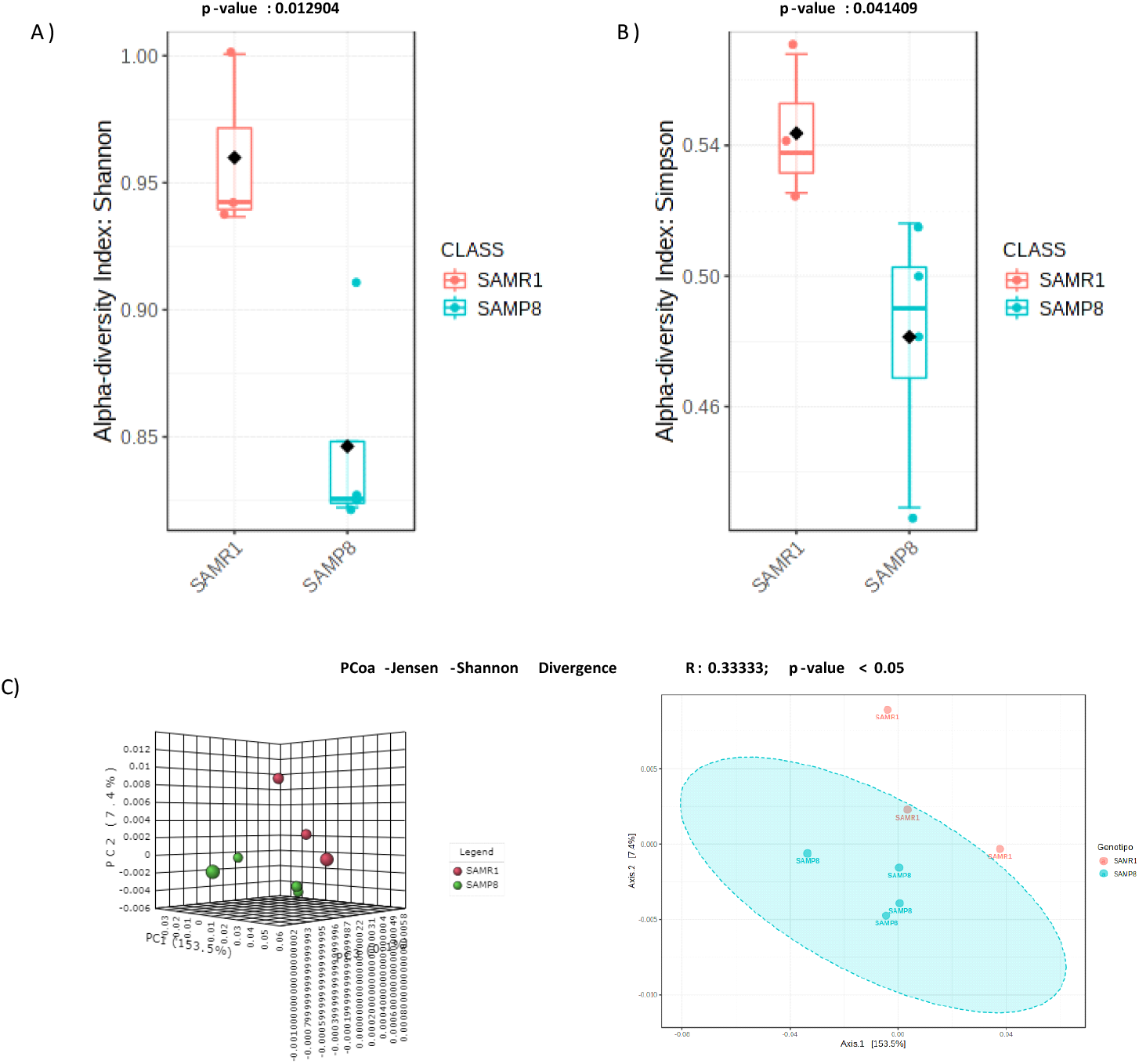
Differential profiles of the gut microbiota between SAMR1 and SAMP8 mice. A) Shannon index (*t*-test, *P* < 0.05). B) Simpson index (*t*-test, *P* < 0.05). C) PCoA analysis of gut bacteria data (Jensen-Shannon Divergence). Analysis of group similarities (ANOSIM).

**Supplementary Figure 6. Effect of DMB treatment in blood glucose and insulin levels.**

A) Blood glucose levels and B) blood insulin levels. ^Δ^ Main effect of treatment, two way ANOVA.

## Funding and disclosure

None of the authors have any actual or potential conflict of interest including any financial, personal or other relationships with other people or organizations that could inappropriately influence their work. This work has been supported by the Spanish Ministry of Economy and Competitiveness (SAF2017-87619-P, Spanish Government).

## Notes

### Competing Interest Statement

The authors have declared no competing interest.

## References

Aguirre V, Uchida T, Yenush L, Davis R, White MF. The c-Jun NH(2)-terminal kinase promotes insulin resistance during association with insulin receptor substrate-1 and phosphorylation of Ser(307). J Biol Chem 2000;275:9047–54.

A.K. Ali, W.A. Banks, V.B. Kumar, Gn Shah, J.L. Lynch, S.A. Farr, M.A. Fleegal-DeMotta, J.E. Morley. Nitric oxide activity and isoenzyme expression in the senescence-accelerated mouse p8 model of Alzheimer’s disease: effects of anti-amyloid antibody and antisense treatments. J. Gerontol. A Biol. Sci. Med. Sci., 64 (2009), pp. 1025–1030

Ballard C, Gauthier S, Corbett A, Brayne C, Aarsland D, Jones E. Alzheimer’s disease. Lancet 2011;377:1019–31.

Banting FG, Best CH. The internal secretion of the pancreas. Lab Clin Med 1922;7:251–66.

Bell R.D., Winkler E.A., Sagare A.P., Singh I., LaRue B., Deane R., Zlokovic B.V, Pericytes control key neurovascular functions and neuronal phenotype in the adult brain and during brain aging, Neuron. 2010; 68(3): 409–427. https://doi.org/10.1016/j.neuron.2010.09.043.

Bell R.D., Winkler E.A., Singh I., Sagare A.P., Deane R., Wu Z., Holtzman D.M., Betsholtz C., Armulik A., Sallstrom J., Berk B.C., Zlokovic B.V., Apolipoprotein E controls cerebrovascular integrity via cyclophilin A, Nature. 2012; 485(7399): 512–516. https://doi.org/10.1038/nature11087.

Biessels GJ, Despa F. Cognitive decline and dementia in diabetes mellitus: mechanisms and clinical implications. Nat Rev Endocrinol. 2018 Oct;14(10):591–604.

Brown MW, Aggleton JP. Recognition memory: what are the roles of the perirhinal cortex and hippocampus? Nat Rev Neurosci. 2001 Jan;2(1):51–61.

Bruning JC, Gautam D, Burks DJ, Gillette J. Schubert M, Orban PC, et al. Role of brain insulin receptor in control of body weight and reproduction. Science 2000;289:2122–5.

Burns, Jeffrey M., Robyn A. Honea, Eric D. Vidoni, Lewis J. Hutfles, William M. Brooks, and Russell H. Swerdlow. 2012. “Insulin Is Differentially Related to Cognitive Decline and Atrophy in Alzheimer’s Disease and Aging.” Biochimica et Biophysica Acta - Molecular Basis of Disease. https://doi.org/10.1016/j.bbadis.2011.06.011.

D.A. Butterfield, H.F. Poon. The senescence-accelerated prone mouse (SAMP8): a model of age-related cognitive decline with relevance to alterations of the gene expression and protein abnormalities in Alzheimer’s disease. Exp. Gerontol., 40 (2005), pp. 774–783.

B. Caballero, I. Vega-Naredo, V. Sierra, C. Huidobro-Fernandez, C. Soria-Valles, D. De Gonzalo-Calvo, D. Tolivia, J. Gutierrez-Cuesta, M. Pallas, A. Camins, M.J. Rodriguez-Colunga, A. Coto-Montes. Favorable effects of a prolonged treatment with melatonin on the level of oxidative damage and neurodegeneration in senescence-accelerated mice. J. Pineal. Res., 45 (2008), pp. 302–311

Cai Z., Qiao P.F., Wan C.Q., Cai M., Zhou N.K., Li Q., Role of Blood-Brain Barrier in Alzheimer’s Disease, J. Alzheimers. Dis. 2018; 63(4): 1223–1234.

A.M. Canudas, J. Gutierrez-Cuesta, M.I. Rodriguez, D. Acuna-Castroviejo, F.X. Sureda, A. Camins, M. Pallas. Hyperphosphorylation of microtubule-associated protein tau in senescence-accelerated mouse (SAM). Mech. Ageing Dev., 126 (2005), pp. 1300–1304

Chen, K.; Zheng, X.; Feng, M.; Li, D.; Zhang, H. Gut Microbiota-Dependent Metabolite Trimethylamine N-Oxide Contributes to Cardiac Dysfunction in Western Diet-Induced Obese Mice. Front Physiol 2017, 8, 139.

Chen, Sifan, Ayana Henderson, Michael C. Petriello, Kymberleigh A. Romano, Mary Gearing, Ji Miao, Mareike Schell, et al. 2019. “Trimethylamine N-Oxide Binds and Activates PERK to Promote Metabolic Dysfunction.” Cell Metabolism 30 (6): 1141-1151.e5. https://doi.org/10.1016/j.cmet.2019.08.021.

Cribbs, D.H., Berchtold, N.C., Perreau, V., Coleman, P.D., Rogers, J., Tenner, A.J., and Cotman, C.W. (2012). Extensive innate immune gene activation accompanies brain aging, increasing vulnerability to cognitive decline and neurodegeneration: a microarray study. J. Neuroinflammation 9, 179.

De Felice FG. Alzheimer’s disease and insulin resistance: translating basic science into clinical applications. J Clin Invest 2013;123:531–9.

De Souza CT, Araujo EP, Bordin S, Ashimine R, Zollner RL, Boschero AC, et al. Consumption of a fat-rich diet activates a proinflammatory response and induces insulin resistance in the hypothalamus. Endocrinology 2005;146:4192–9.

Erny D, Hrabě de Angelis AL, Jaitin D, Wieghofer P, Staszewski O, David E, Keren-Shaul H, Mahlakoiv T, Jakobshagen K, Buch T, Schwierzeck V, Utermöhlen O, Chun E, Garrett WS, McCoy KD, Diefenbach A, Staeheli P, Stecher B, Amit I, Prinz M. Host microbiota constantly control maturation and function of microglia in the CNS. Nat Neurosci. 2015 Jul;18(7):965–77.

Fang, Chuo, Hang Xu, Shaodong Guo, Susanne U. Mertens-Talcott, and Yuxiang Sun. 2018. “Ghrelin Signaling in Immunometabolism and Inflamm-Aging.” In Advances in Experimental Medicine and Biology. https://doi.org/10.1007/978-981-13-1286-1_9.

Fernandes ML, Saad MJ, Velloso LA. Effects of age on elements of insulin-signaling pathway in central nervous system of rats. Endocrine 2001;16:227–34.

Fernandez AM, Torres-Aleman I. The many faces of insulin-like peptide signalling in the brain. Nat Rev Neurosci 2012;13:225–39.

Ferreira ST, Clarke JR, Bomfim TR, De Felice FG. Inflammation, defective insulin signaling, and neuronal dysfunction in Alzheimer’s disease. Alzheimers Dement. 2014 Feb;10(1 Suppl):S76–83.

Frazier, Hilaree N., Adam O. Ghoweri, Katie L. Anderson, Ruei Lung Lin, Nada M. Porter, and Olivier Thibault. 2019. “Broadening the Definition of Brain Insulin Resistance in Aging and Alzheimer’s Disease.” Experimental Neurology 313 (October 2018): 79–87. https://doi.org/10.1016/j.expneurol.2018.12.007.

Frisardi V, Solfrizzi V, Seripa D, Capurso C, Santamato A, Sancarlo D. Metabolic-cognitive syndrome: a cross-talk between metabolic syndrome and Alzheimer’s disease. Ageing Res Rev 2010;9:399–417.

Frolich L, Blum-Degen D, Bernstein HG, Engelsberger S, Humrich J, Laufer S. Brain insulin and insulin receptors in aging and sporadic Alzheimer’s disease. J Neural Transm 1998;105:423–38.

A. Fukunari, A. Kato, Y. Sasaki, T. Yoshimoto, S. Ishiura, K. Suzuki, T. Nakajima. Colocalization of prolyl endopeptidase and amyloid β-peptide in brains of senescence-accelerated mouse. Neurosci. Lett., 176 (1994), pp. 201–204

Gao, Xiang, Jie Xu, Chengzi Jiang, Yi Zhang, Yong Xue, Zhaojie Li, Jingfeng Wang, Changhu Xue, and Yuming Wang. 2015. “Fish Oil Ameliorates Trimethylamine N-Oxide-Exacerbated Glucose Intolerance in High-Fat Diet-Fed Mice.” Food and Function 6 (4): 1117–25. https://doi.org/10.1039/c5fo00007f.

Gregory JC, Buffa JA, Org E, Wang Z, Levison BS, Zhu W, Wagner MA, Bennett BJ, Li L, DiDonato JA, Lusis AJ, Hazen SL. Transmission of atherosclerosis susceptibility with gut microbial transplantation. J Biol Chem. 2015 27;290(9):5647–60.

Hotamisligil GS, Shargill NS, Spiegelman BM. Adipose expression of tumor necrosis factor-alpha: direct role in obesity-linked insulin resistance. Science. 1993 Jan 1;259(5091):87–91.

Janeiro, Manuel H., María J. Ramírez Fermin I. Milagro, J. Alfredo Martínez, and Maite Solas. 2018. “Implication of Trimethylamine N-Oxide (TMAO) in Disease: Potential Biomarker or New Therapeutic Target.” Nutrients 10 (10). https://doi.org/10.3390/nu10101398.

Janeiro, Manuel H., María J. Ramírez, and Maite Solas. 2021. “Dysbiosis and Alzheimer’s Disease: Cause or Treatment Opportunity?” Cellular and Molecular Neurobiology, no. 0123456789. https://doi.org/10.1007/s10571-020-01024-9.

Ke, Yilang, Dang Li, Mingming Zhao, Changjie Liu, Jia Liu, Aiping Zeng, Xiaoyun Shi, et al. 2018. “Gut Flora-Dependent Metabolite Trimethylamine-N-Oxide Accelerates Endothelial Cell Senescence and Vascular Aging through Oxidative Stress.” Free Radical Biology and Medicine 116 (January): 88–100. https://doi.org/10.1016/j.freeradbiomed.2018.01.007.

Y. Kitamura, X.H. Zhao, T. Ohnuki, M. Takei, Y. Nomura. Age-related changes in transmitter glutamate and NMDA receptor/channels in the brain of senescence-accelerated mouse. Neurosci. Lett., 137 (1992), pp. 169–172

Koeth RA, Wang Z, Levison BS, Buffa JA, Org E, Sheehy BT, Britt EB, Fu X, Wu Y, Li L, Smith JD, DiDonato JA, Chen J, Li H, Wu GD, Lewis JD, Warrier M, Brown JM, Krauss RM, Tang WH, Bushman FD, Lusis AJ, Hazen SL. Intestinal microbiota metabolism of L-carnitine, a nutrient in red meat, promotes atherosclerosis. Nat Med. 2013;19(5):576–85.

Konner AC, Bruning JC. Toll-like receptors: linking inflammation to metabolism. Trends Endocrinol Metab 2011;22:16–23

V.B. Kumar, M. Franko, W.A. Banks, P. Kasinadhuni, S.A. Farr, K. Vyas, V. Choudhuri, J.E. Morley. Increase in presenilin 1 (PS1) levels in senescence-accelerated mice (SAMP8) may indirectly impair memory by affecting amyloid precursor protein (APP) processing. J. Exp. Biol., 212 (Pt 4) (2009), pp. 494–498

Kushner JA. The role of aging upon beta cell turnover. J Clin Invest 2013;123:990–5.

Kuusisto, Johanna, Keijo Koivisto, Leena Mykkänen, Eeva Liisa Helkala, Matti Vanhanen, Tuomo Hänninen, Kalevi Pyörälä, Paavo Riekkinen, and Markku Laakso. 1993. “Essential Hypertension and Cognitive Function: The Role of Hyperinsulinemia.” Hypertension. https://doi.org/10.1161/01.HYP.22.5.771.

Lee CK, Weindruch R, Prolla TA. Gene-expression profile of the ageing brain in mice. Nat Genet 2000;25:294–7.

Mokdad AH, Ford ES, Bowman BA,, Dietz WH, Vinicor F, Bales VS. Prevalence of obesity, diabetes, and obesity-related health risk factors, 2001; JAMA 2003;289:76–9.

J.E. Morley, V.B. Kumar, A.E. Bernardo, S.A. Farr, K. Uezu, N. Tamosa, J.F. Flood. Beta-amyloid precursor polypeptide in SAMP8 mice affects learning and memory. Peptides, 21 (2000), pp. 1761–1767.

Y. Nomura, Y. Kitamura, T. Ohnuki, T. Arima, Y. Yamanaka, K. Sasaki, Y. Oomura. Alterations in aceylcholine, NMDA, benzodiazepine receptors and protein kinase C in the brain of the senescence-accelerated mouse: an animal model useful for studies on cognitive enhancers. Behav. Brain Res., 83 (1997), pp. 51–55

Norden, D.M., and Godbout, J.P. (2013). Review: microglia of the aged brain: primed to be activated and resistant to regulation. Neuropathol. Appl. Neurobiol. 39, 19–34.

O’Connor A, Quizon PM, Albright JE, Lin FT, Bennett BJ. Responsiveness of cardiometabolic-related microbiota to diet is influenced by host genetics. Mamm Genome. 2014 Dec;25(11-12):583–99.

Oellgaard, Jens, Signe Abitz Winther, Tobias Schmidt Hansen, Peter Rossing, and Bernt Johan von Scholten. 2017. “Trimethylamine N-Oxide (TMAO) as a New Potential Therapeutic Target for Insulin Resistance and Cancer.” Current Pharmaceutical Design 23 (25): 3699–3712. https://doi.org/10.2174/1381612823666170622095324.

O’Toole, Paul W., and Ian B. Jeffery. 2015. “Gut Microbiota and Aging.” Science 350 (6265): 1214–15. https://doi.org/10.1126/science.aac8469.

Pedersen, M., Bruunsgaard, H., Weis, N., Hendel, H. W., Andreassen, B. U., Eldrup, E., et al. (2003). Circulating levels of TNF-alpha and IL-6-relation to truncal fat mass and muscle mass in healthy elderly individuals and in patients with type-2 diabetes. Mech. Ageing Dev. 124, 495–502.

Posey KA, Clegg DJ, Printz RL, Byun J, Morton GJ, Vivekanandan-Giri A, et al. Hypothalamic proinflammatory lipid accumulation, inflammation, and insulin resistance in rats fed a high-fat diet. Am J Physiol Endocrinol Metab 2009;296:E1003–12.

Rothhammer V, Mascanfroni ID, Bunse L, Takenaka MC, Kenison JE, Mayo L, Chao CC, Patel B, Yan R, Blain M, Alvarez JI, Kébir H, Anandasabapathy N, Izquierdo G, Jung S, Obholzer N, Pochet N, Clish CB, Prinz M, Prat A, Antel J, Quintana FJ. Type I interferons and microbial metabolites of tryptophan modulate astrocyte activity and central nervous system inflammation via the aryl hydrocarbon receptor. Nat Med. 2016 Jun;22(6):586–97.

Rui L, Aguirre V, Kim JK, Shulman GI, Lee A, Corbould A, Dunaif A, White MF. Insulin/IGF-1 and TNF-alpha stimulate phosphorylation of IRS-1 at inhibitory Ser307 via distinct pathways. J Clin Invest. 2001 Jan;107(2):181–9.

Solas M, Aisa B, Mugueta MC, Del Río J, Tordera RM, Ramírez MJ. Interactions between age, stress and insulin on cognition: implications for Alzheimer’s disease. Neuropsychopharmacology. 2010 Jul;35(8):1664–73.

Stolk, Ronald P., Monique M.B. Breteler, Alewijn Ott, Huibert A.P. Pols, Steven W.J. Lamberts, Diederick E. Grobbee, and Albert Hofman. 1997. “Insulin and Cognitive Function in an Elderly Population the Rotterdam Study.” Diabetes Care. https://doi.org/10.2337/diacare.20.5.792.

R. Strong, V. Reddy, J.E. Morley. Cholinergic deficits in the septal–hippocampal pathway of the SAM-P/8 senescence accelerated mouse. Brain Res., 966 (2003), pp. 150–156.

M. Takemura, S. Nakamura, I. Akiguchi, M. Ueno, N. Oka, S. Ishikawa, A. Shimada, J. Kimura, T. Takeda. Beta/A4 protein-like immunoreactive granular structures in the brain of senescence-accelerated mouse. Am. J. Pathol., 142 (1993), pp. 1887–1897.

Thaler JP, Guyenet SJ, Dorfman MD, Wisse BE, Schwartz MW. Hypothalamic inflammation: marker or mechanism of obesity pathogenesis? Diabetes 2013;62:2629–34.

Velloso LA, Schwartz MW. Altered hypothalamic function in diet-induced obesity. Int J Obes (Lond) 2011;35:1455–65.

Vogt MC, Bruning JC. CNS insulin signaling in the control of energy homeostasis and glucose metabolism: from embryo to old age. Trends Endocrinol Metab 2013;24:76–84.

Wang, Zeneng, Adam B Roberts, Jennifer A Buffa, Bruce S Levison, Weifei Zhu, Elin Org, Xiaodong Gu, et al. 2015. “Non-Lethal Inhibition of Gut Microbial Trimethylamine Production for the Treatment of Atherosclerosis Assisted in Gene Cloning, Protein Purification, Choline Transport and TMA Lyase Activity Assay HHS Public Access.” Cell 163 (7): 1585–95. https://doi.org/10.1016/j.cell.2015.11.055.

Weiss S, Xu Z, Amir A, Peddada S, Bittinger K, Gonzalez A, et al. Effects of library size variance, sparsity, and compositionality on the analysis of microbiome data. PeerJ Prepr. 2015.

Winkler E.A., Sengillo J.D., Bell R.D., Wang J., Zlokovic B.V., Blood-spinal cord barrier pericyte reductions contribute to increased capillary permeability, J. Cereb. Blood. Flow. Metab. 2012; 32(10): 1841–1852. https://doi.org/10.1038/jcbfm.2012.113.

Winkler E.A., Sengillo J.D., Sullivan J.S., Henkel J.S., Appel S.H., Zlokovic B.V., Blood-spinal cord barrier breakdown and pericyte reductions in amyotrophic lateral sclerosis, Acta. Neuropathol. 2013; 125(1): 111–120. https://doi.org/10.1007/s00401-012-1039-8.

Zhang G, Li J, Purkayastha S, Tang Y, Zhang H, Yin Y, et al. Hypothalamic programming of systemic ageing involving IKK-beta, NF-kappaB and GnRH. Nature 2013;497:211–6.

Zhang X, Zhang G, Zhang H, Karin M, Bai H, Cai D. Hypothalamic IKKbeta/NF-kappaB and ER stress link overnutrition to energy imbalance and obesity. Cell 2008;135:61–73.

Zhao WQ, Chen H, Quon MJ, Alkon DL. Insulin and the insulin receptor in experimental models of learning and memory. Eur J Pharmacol 2004;490:71–81.

